# Numerosity tuning in human association cortices and local image contrast representations in early visual cortex

**DOI:** 10.1101/2021.03.28.437364

**Authors:** Jacob M. Paul, Martijn van Ackooij, Tuomas C. ten Cate, Ben M. Harvey

## Abstract

Human early visual cortex response amplitudes monotonically increase with numerosity (object number), regardless of object size and spacing. However, numerosity is typically considered a high-level visual or cognitive feature, while early visual responses follow image contrast in the spatial frequency domain. We found that, at fixed contrast, aggregate Fourier power (at all orientations and spatial frequencies) followed numerosity closely but nonlinearly with little effect of object size, spacing or shape. This would allow straightforward numerosity estimation from spatial frequency domain image representations. Using 7T fMRI, we showed monotonic responses originate in primary visual cortex (V1) at the stimulus’s retinotopic location. Responses here and in neural network models followed aggregate Fourier power more closely than numerosity. Truly numerosity tuned responses emerged after lateral occipital cortex and were independent of retinotopic location. We propose numerosity’s straightforward perception and neural responses may have built on behaviorally beneficial spatial frequency analyses in simpler animals.

## Introduction

Humans and many other animals use visual numerosity, the number of items in a set, to guide behavior. Many species have neurons tuned to numerosity, decreasing in response amplitude with distance from a specific preferred numerosity (1–3). Functional magnetic resonance imaging (fMRI) has identified numerosity-tuned neuronal populations in specific areas of human association cortex using both population receptive field modeling approaches (4, 5) and fMRI adaptation (or repetition suppression)(6). Other fMRI studies using multivoxel pattern analyses (7, 8) and representational similarity analyses (9) also support the existence of numerosity-tuned neural populations in the human brain. Response properties of these neurons mirror properties of numerosity perception (3, 6, 10). Numerosity perception is correlated with numerosity tuned responses between trials (3, 10), and repetition suppression (6) and multivoxel pattern discriminability between individuals (11, 12).

There remains considerable debate over how such numerosity-tuned responses are derived from visual inputs. One view proposes that numerosity tuning and perception reflect non-numerical image features that are often correlated with numerosity, like density (13), or contrast energy at high spatial frequencies (14). However, growing convergent evidence from psychophysical, neuroimaging and computational research indicates numerosity itself is represented and perceived (15–19). These views could be reconciled by a non-numerical image feature from which numerosity could be estimated regardless of other image parameters like item size and spacing.

Computational modeling shows numerosity-tuned responses in various neural networks that are not trained for numerosity discrimination. The first of these models (20) pre-dates the discovery of numerosity tuned neurons. It implements specific stages that: (1) detect where contrast lies in the image; (2) normalize the local contrast so that each item contributes equally; (3) sum normalized contrast to give a monotonically increasing and decreasing response to numerosity; (4) weight these monotonic responses differently to give numerosity-tuned responses with different numerosity preferences. A simple unsupervised network can develop monotonically increasing and tuned responses with no need for monotonically decreasing responses to numerosity (as inhibitory synapses are sufficient) (21). Neither model shows how the monotonic stage disregards size and spacing. It has since been shown that monotonic (22) and tuned (23) responses to numerosity emerge in a probabilistic hierarchical generative network trained only to efficiently encode the image and maximize the likelihood of reconstructing the image, and even in a randomly-weighted network. In this model, the first stage decomposes the image using spatial receptive fields with surround suppression, as in the early visual system. The resulting monotonic responses to numerosity are spatially selective (22), but responses are almost invariant to item size and spacing without the need for explicit object individuation or size normalization (19). Another class of neural network model, deep convolutional neural networks, also show monotonic and numerosity-tuned units, even in randomly-weighted networks (24). Here monotonic units emerge early in the network and feed into numerosity-tuned units, where different weights on these inputs give numerosity-tuned responses with different numerosity preferences.

EEG and fMRI results show that early visual cortex responses to numerosity stimuli appear to monotonically increase with numerosity, regardless of item size or spacing (25, 26). This monotonic response emerges very quickly after stimulus presentation, suggesting it reflects feedforward processing, and is not computed from other parameters like stimulus area and density. This early visual cortex response to numerosity is surprising, as numerosity is generally considered a relatively high-level visual feature. Early visual neurons respond to image contrast in the frequency domain: at specific positions, orientations and spatial frequencies (27). So, it is unclear how early visual neural responses could closely follow numerosity regardless of size and spacing, although such responses spontaneously emerge somehow in the image representations of computational models.

Association cortex recording sites with numerosity-tuned responses largely overlap with higher visual field maps (5), but their spatial population receptive fields (pRF) do not necessarily overlap with the numerosity stimulus area (5, 28), and numerosity preferences are unrelated to pRF position or size (28). Numerosity tuning in functionally homologous macaque brain areas locations (29) also does not require the responding single neurons to have spatial receptive fields including the stimulus region, or even have discernible receptive fields at all (30). Indeed, spatial receptive field properties of numerosity-tuned neurons do not influence numerosity preferences, tuning width or firing rate. For early visual monotonic responses, some properties of EEG event related potentials suggest these emerge in V2 or V3 (31), but their precise visual field map and retinotopic location remains unclear.

Here we use computational modeling of human 7T fMRI data to ask precisely where in the early visual cortex monotonic responses emerge, whether they can be explained by the spatial frequency domain image representation in the early visual cortex, and how they relate to location-independent numerosity tuned responses in the human association cortices.

## Results

### Numerosity response profiles differ between early visual cortex and association cortices

We presented fixed contrast displays of gradually changing numerosity to our participants while collecting 7T fMRI data (see Methods). We included different stimulus configurations that held total item area, individual item size, or total item perimeter constant across numerosities, or placed items in a dense group (Fig. 1). These configurations varied item size and spacing considerably but produced similar responses.

**Fig. 1:**
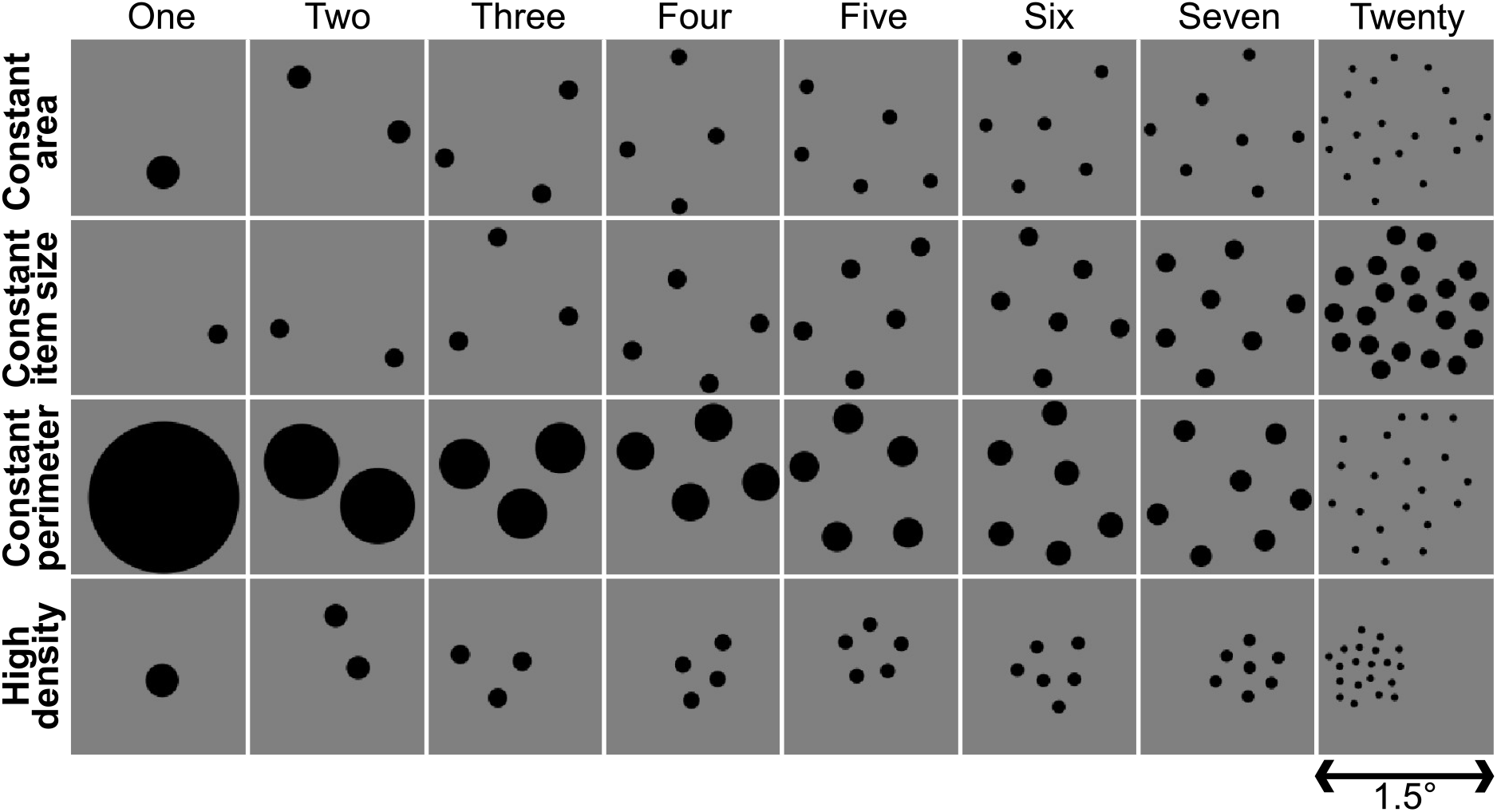
Example displays from each stimulus configuration and numerosity. Different cortical locations showed different relationships between numerosity and response amplitude. These were well captured by response models with monotonically increasing, monotonically decreasing, or tuned responses to log(numerosity) at different locations (Supplementary Fig. 2c–d). We compared the response variance explained by these models in cross-validated data (Fig. 2a and Supplementary Fig. 4). Separate visual field mapping data demonstrate that monotonically increasing responses were consistently found only in early visual cortex’s central visual field representation. Numerosity tuned response were found outside early visual cortex, in previously described areas of temporal-occipital, parietal-occipital, superior parietal, and frontal cortices containing topographic numerosity maps (5). These largely overlapped with higher extrastriate visual field maps. Monotonically decreasing responses were found next to areas showing tuned responses.

### Early visual monotonic responses

Early visual (V1–V3, LO1) responses were consistently predicted more closely by monotonically increasing rather than tuned responses to numerosity (Fig. 2b). Critically, model fits depended on the recording sites’ visual field position preferences: those near fixation (the stimulus location) showed better fits, gradually decreasing to zero into the periphery. These progressions were well captured by cumulative Gaussian sigmoid functions (Supplementary Fig. 5c). Inflection points of these sigmoid curves fell at eccentricities between one and two degrees of visual angle (Fig. 2b and Supplementary Fig. 2a).

**Fig. 2:**
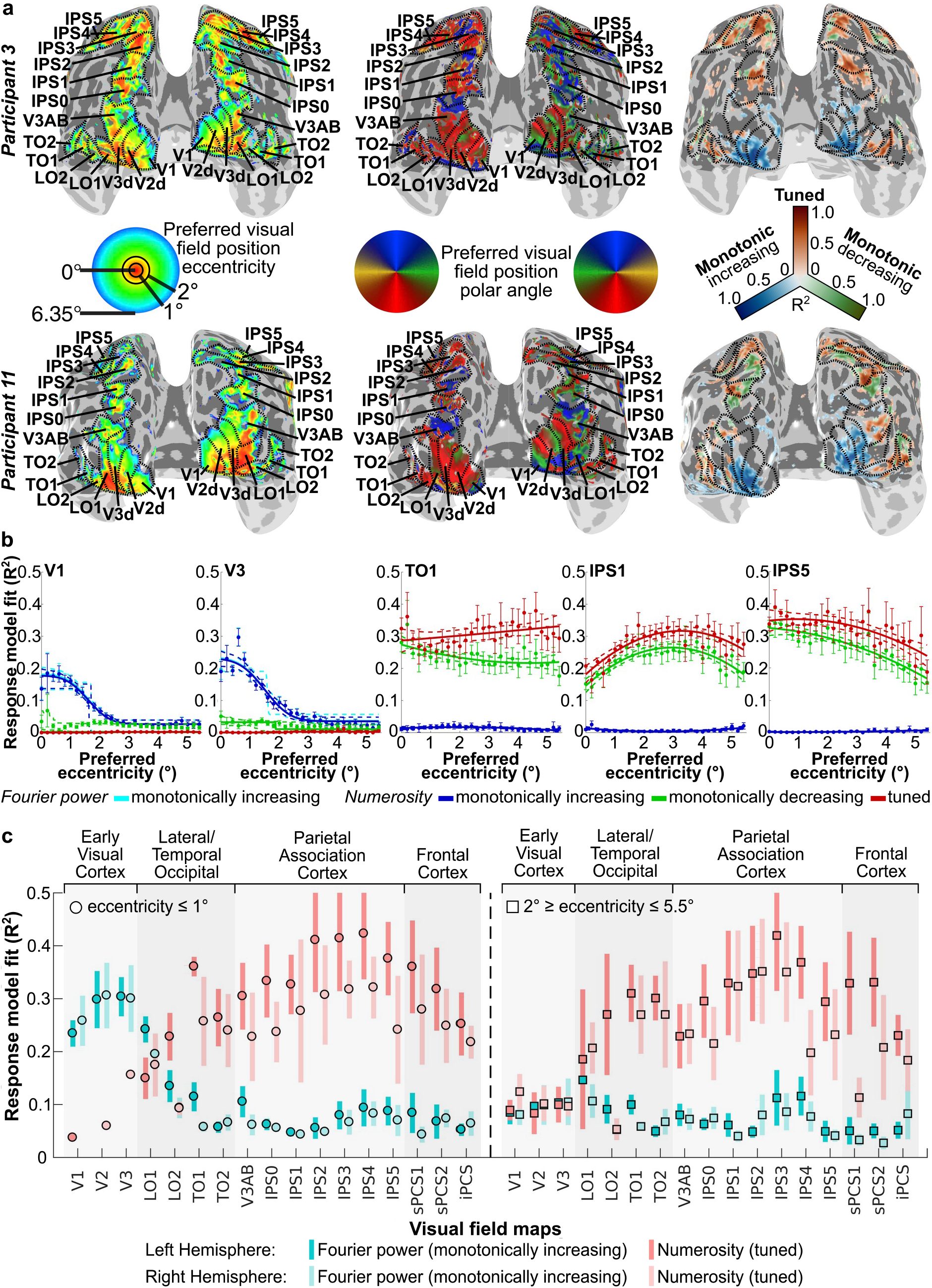
Relationships between responses to numerosity and visual field position. **a** Visual position preferences (eccentricity and polar angle, left and middle) and best fitting numerosity model (right, colors) at each cortical location, for two illustrative participants. Dashed black lines and labels show visual field map borders and names respectively. **b** Progression of each numerosity model’s fit with preferred visual field eccentricity (colors) in representative visual field maps (grouped across participants). Cyan shows fits of monotonically increasing responses to aggregate Fourier power, which are often hidden by the very similar blue line (monotonically increasing responses to numerosity) that is drawn on top. Filled circles show mean variance explained per eccentricity bin, error bars show the standard error of the mean. Solid lines show the best fit to changes with eccentricity, dashed lines are bootstrap 95% confidence intervals determined by bootstrapping. **c** Model fit variance explained by numerosity (tuned, reds) and aggregate Fourier power (monotonically increasing, blues) response models for each visual field map hemisphere at eccentricities below 1° (left) and between 2° and 5.5° (right). Points represent the population marginal mean, error bars are 95% confidence intervals; non-overlapping error bars show significant differences at *p* < 0.05.

A location-specific monotonic response to numerosity that has emerged by V1 is perhaps surprising: numerosity is generally considered a complex visual or cognitive feature, and responses of V1 neurons depend on contrast at specific orientations and spatial frequencies within their receptive field. Images are often transformed into this spatial frequency domain using a 2-dimensional Fourier decomposition, which similarly separates changes across an image into the contrast at each orientation and spatial frequency (32). We therefore reasoned that aggregate Fourier power (across all orientations and spatial frequencies) in the spatial frequency domain might closely predict the measured aggregate response of V1 populations. We summed the absolute Fourier decomposition of the displays within the first spatial frequency harmonic, across all orientations (Fig. 3a–b). This revealed that aggregate Fourier power followed numerosity closely, but nonlinearly, and similarly in all stimulus configurations (Fig. 3c). The aggregate power of the second harmonic showed a similar pattern at half the amplitude. Indeed, aggregate Fourier power changed very little over a wide range of item sizes, spacings and shapes that were not tested in our experiments. Fourier power followed numerosity closely except in extreme cases, and except for and a slight increase in Fourier power with item spacing (Fig. 3d–g and Supplementary Figs. 7–10). Global changes in item contrast do not affect perceived numerosity until items become less visible (33). However, Fourier power increased linearly with the contrast between items and the background (Fig. 3h). This effect could therefore be compensated for simply by divisive normalization: dividing the Fourier power by the mean contrast of the items. Using a mixture of black and white items gave very slightly (approximately 1.6%) higher Fourier power than using black items only, which is unsurprising because the difference in luminance between any item and the rest of the display increased slightly here.

**Fig. 3:**
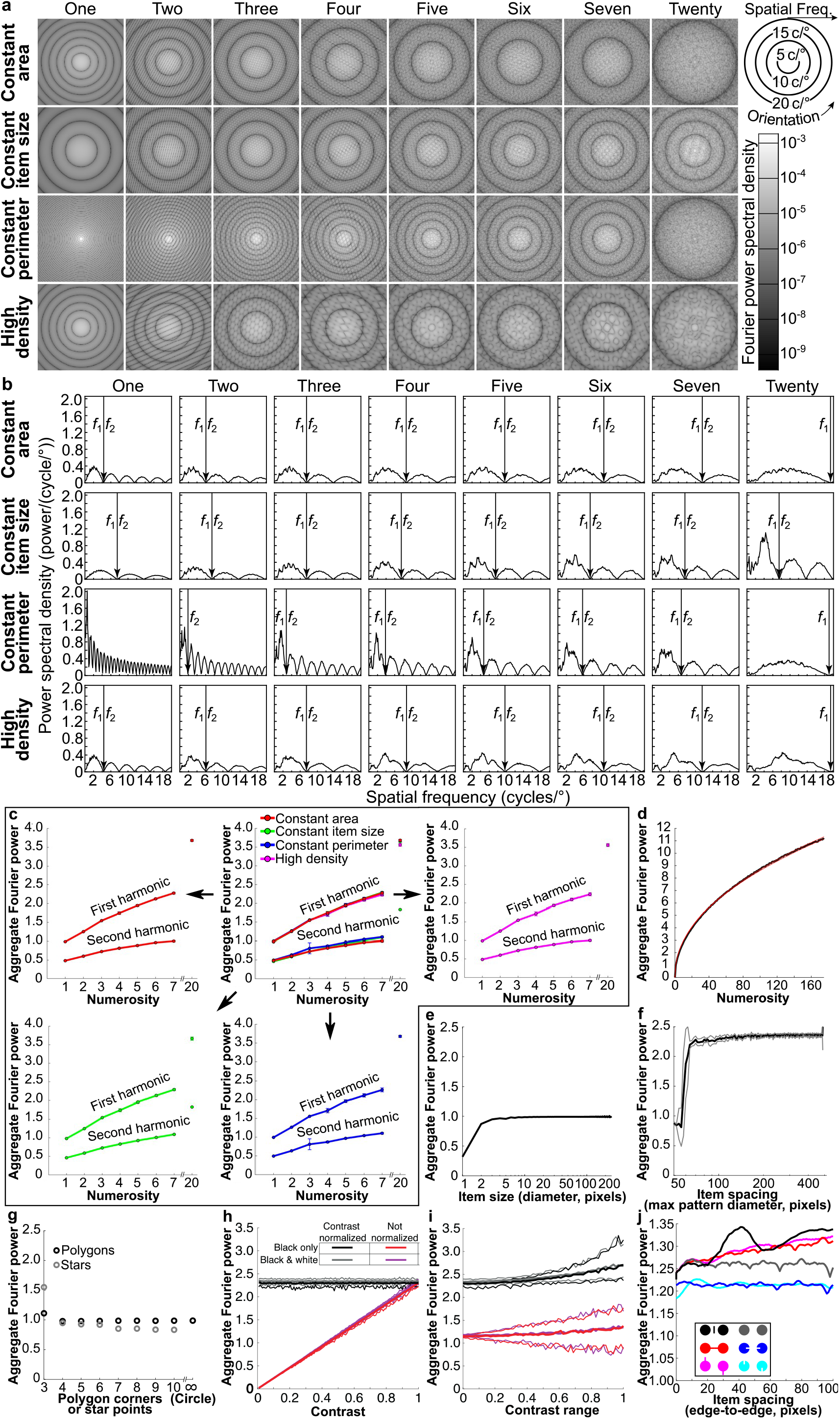
Aggregate Fourier power followed numerosity closely with little effect of item size, spacing, shape or connectedness. **a** 2-dimensional Fourier transforms for each image in Fig. 1. **b** Fourier power spectral density at each spatial frequency, collapsed over orientation. The limit of the first harmonic (*f*1) was used to determine the aggregate Fourier power of each image. **c** Aggregate Fourier power **b** follows numerosity closely and similarly across stimulus configurations. We divided aggregate Fourier power by the product of number display pixels and the square root of two here, making the power of one circle approximately one for any display resolution. Points show the mean power of all stimuli of the same numerosity in the same stimulus configuration, error bars show the standard deviation. Upper middle panel shows all configurations overlaid; other panels show individual configurations. **d** Aggregate Fourier power as a function of numerosity for a fixed item size (black line). Red lines show *y* = *x*^0.466^ and *y* = *x*^0.470^, which approximate the observed relationship. **e** Aggregate Fourier power of one circle of different diameters was approximately constant for diameters above 3 pixels: smaller circles were inaccurately rendered. **f** Aggregate Fourier power for a group of 7 circles (each 16 pixels diameter) increased slightly with spacing for stimulus area diameters above 70 pixels. Smaller areas had little space between items: at 50 pixels all items touch (Supplementary Fig. 9). Gray lines show the standard deviation across displays. **g** Aggregate Fourier power was approximately constant for regular polygons above 3 corners. Triangles and 3-pointed stars had greater Fourier power than circles, while other stars had less Fourier power. **h** Aggregate Fourier power for a group of 7 circles (each 16 pixels diameter) increased linearly as the absolute Weber contrast of items increased (red). Therefore, normalizing Fourier power by dividing by item contrast gave an approximately constant value for all contrasts (black). Fourier power was slightly higher for a random mixture of black and white items (magenta and gray) than only black items (red and black). Fine lines show the 95% confidence intervals across displays. (I) When item contrasts were randomly chosen from a range centered on 0.5 contrast, the mean Fourier power across displays increased only slightly as this range increased. The 95% confidence intervals increased considerably. **j** Effects of connections by a bar and an illusory contour on Fourier power, together with control dot pairs with the same change to the dot but no connection and no reduction in perceived numerosity. Compared to dots alone, bars increased Fourier power while illusory contour inducers reduced Fourier power, regardless of connectedness.

Numerosity perception is less accurate in displays where different items have variable contrasts than in fixed-contrast displays (34). As we increased the range of item contrasts in a display, the range of Fourier powers in different displays also increased considerably (Fig. 3i): the 95% confidence intervals were approximately 300% greater when the range of contrasts was 0.85 than when it was 0.1, regardless of whether Fourier power is normalized by the mean contrast of the items. The mean Fourier power also increased, but only by approximately 10%.

Connecting dot pairs with bars reduced the numerosity of items perceived in the display. This perceived numerosity was originally compared to numerosity perceived when bars were placed among the dots but fully unconnected (14, 35)(Fig. 3j, black inset). The Fourier power of a connected pair (red) was lower than fully unconnected dots and bars (black), superficially predicting the connectedness illusion. However, rotating the dots and bars to break the connection (magenta) had the same lower Fourier power while it is perceived as more numerous. Furthermore, the Fourier power of the connected pair was greater than the dots alone (gray), while connected pairs are perceived as less numerous than dots alone. Connecting illusory contour inducers (blue) also reduce perceived numerosity and had lower Fourier power than dots alone, but these inducers again had the same effect when rotated to break the connection (cyan). Therefore, while bars and illusory contour affected Fourier power, these effects were not consistent with perceptual effects: the connectedness illusion likely results from higher-order segmentation processes (36).

Several other non-numerical stimulus features have been proposed to account for numerosity perception because they are correlated with numerosity in some cases, though none shows this generalization across displays. We quantified several such features for each numerosity and stimulus configurations (17, 18) and tested predictions of response models that monotonically follow these features against the recorded early visual (V1-V3) responses. We tested predictions of position selective pRFs responding either to dot bodies (i.e. luminance) or edges (i.e. contrast) (5). All of these models predicted response amplitudes significantly less well than log(numerosity) (all *Z* ≥ 3.21, *p* < .0013 in paired Wilcoxon signed rank tests, false discovery rate (FDR) corrected (37) for all comparisons against the log(numerosity) response model, *n*=20 hemisphere measurements) (Fig. 4a).

**Fig. 4:**
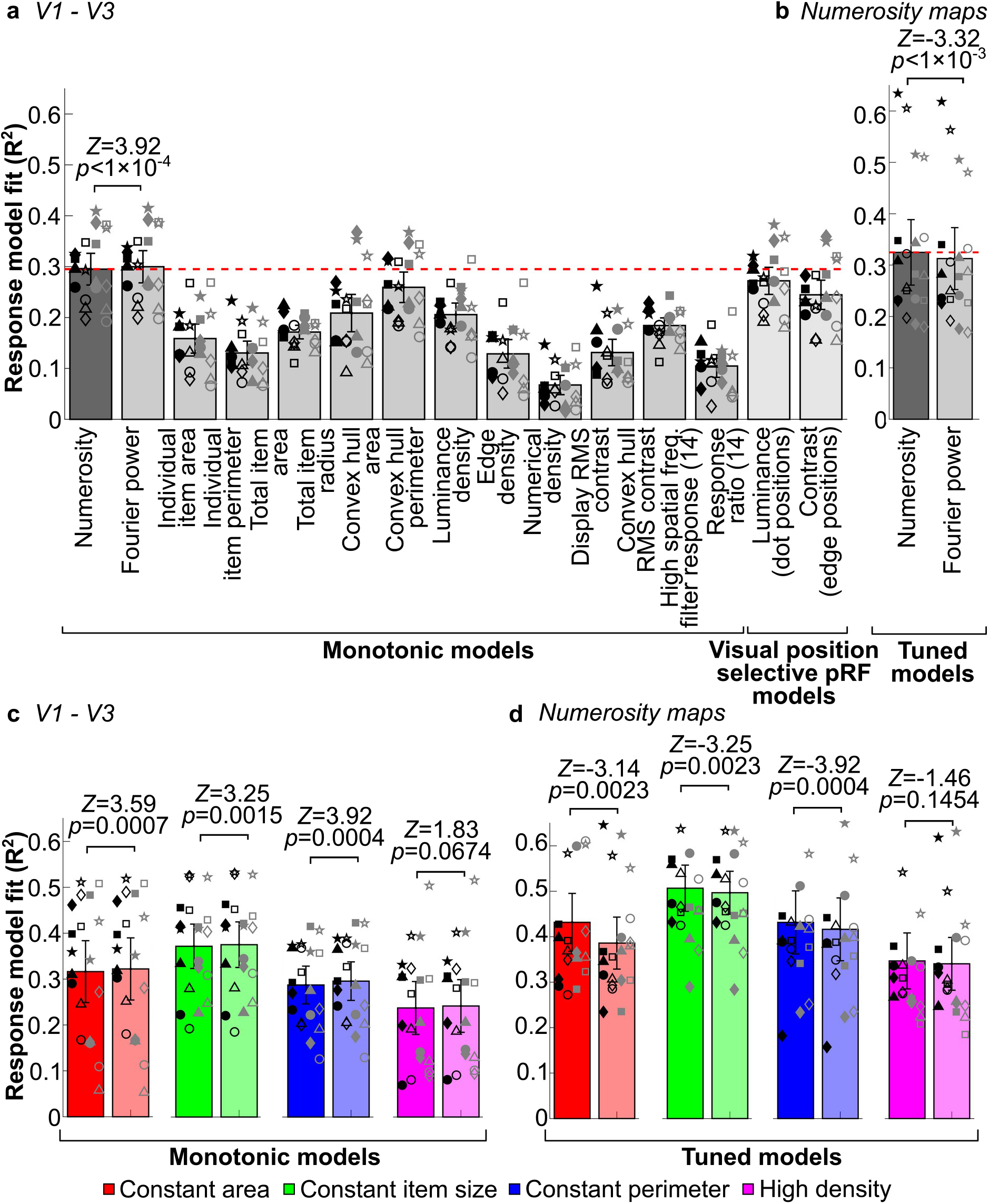
Responses to aggregate Fourier power predict early visual monotonic responses better than numerosity, but numerosity predicts tuned responses better. **a** Variance explained by mono response models for numerosity and non-numerical visual features, and visual position selective pRF models in V1-V3. **b** Variance explained by tuned response models for numerosity and aggregate Fourier power in the association cortex numerosity maps. **c** Variance explained by monotonic response models for numerosity (saturated bars) and Fourier power (unsaturated bars) in V1-V3 in each stimulus configuration. **d** Variance explained by tuned response models for numerosity and Fourier power in the association cortex numerosity maps in each stimulus configuration. Bar height is the mean variance explained across all maps. Error bars show 95% confidence intervals, reflecting the range of fits between individual maps. Markers show the median variance explained for each measure: different shapes are different participants; filled and unfilled symbols are odd and even runs; black is left hemisphere and gray is right hemisphere. P-values show significance of differences in paired Wilcoxon signed rank tests.

However, early visual responses were significantly better predicted by monotonically increasing responses to log(Fourier power) than log(numerosity) (and all other models), both across all stimulus configurations (Median difference = 0.0054, *Z* = 3.92, *p* = 0.000095, FDR corrected with the other comparisons against the log(numerosity) model) and within each configuration, though this difference is not significant in the high density configuration (*Z* = 1.83, *p* = 0.0674, FDR corrected for the comparisons in different stimulus configurations) (Fig. 4c). Conversely, numerosity tuned models predict the tuned response of six previously identified numerosity maps (5) significantly better than models tuned to aggregate Fourier power across all stimulus configurations (Median difference = 0.0104, *Z* = -3.32, *p* = 0.00089) (Fig. 4b), as previously shown for other non-numerical features (17, 18). Numerosity-tuned models also fit better within each stimulus configuration, though this difference is not significant in the high density configuration (*Z* = -1.46, *p* = 0.1454, FDR corrected for the comparisons in different stimulus configurations) (Fig. 4d).

### Comparison to neural network models

Although it is beyond the scope of this study to implement neural network models and test how they respond to our stimuli, the studies of Stoianov and Zorzi (22) (their Supplementary Figure 4A) and Kim and colleagues (24) (their Figure 5B) show that the monotonic responses in their networks increase nonlinearly with numerosity. We therefore ask whether this nonlinearity follows the nonlinear relationship between numerosity and Fourier power. We took the data from 8 lines shown in each of these studies and normalized these to fall within the same range (Fig. 5). This revealed that both studies show very similar relationships between numerosity and response amplitude, which was not apparent in the original figures because they use log (Figure 5A) and linear (Figure 5B) numerosity axes respectively. We then rescaled the log(numerosity) function and the relationship between numerosity and aggregate Fourier power to best fit all these data points. Finally, we fit the quadratic function that best follows these data points (*response* ∝ -0.0147 × *numerosity*^2^ + *numerosity*). This set of lines was better correlated with aggregate Fourier power than log(numerosity) (*p* = 0.000001, *t* = 7.62, *n* = 16 lines, in a paired t-test of correlation coefficients). There was no significant difference in the correlation of these lines with aggregate Fourier power and the best fitting quadratic function (*p* = 0.53, *t* = 0.63, *n* = 16). Therefore, like the responses of the early visual cortex, the monotonic responses in a hierarchical generative network trained to efficiently encode images (22) and in an untrained deep convolutional neural network (24) both follow aggregate Fourier power more closely than numerosity. We did not extend this comparison to numerosity-tuned responses, which exhibit a range of preferred numerosities rather than having a single shape. Furthermore, the population tuning functions of our fMRI voxels are broader and not straightforwardly comparable to the single-unit tuning functions of neural network models.

**Fig. 5:**
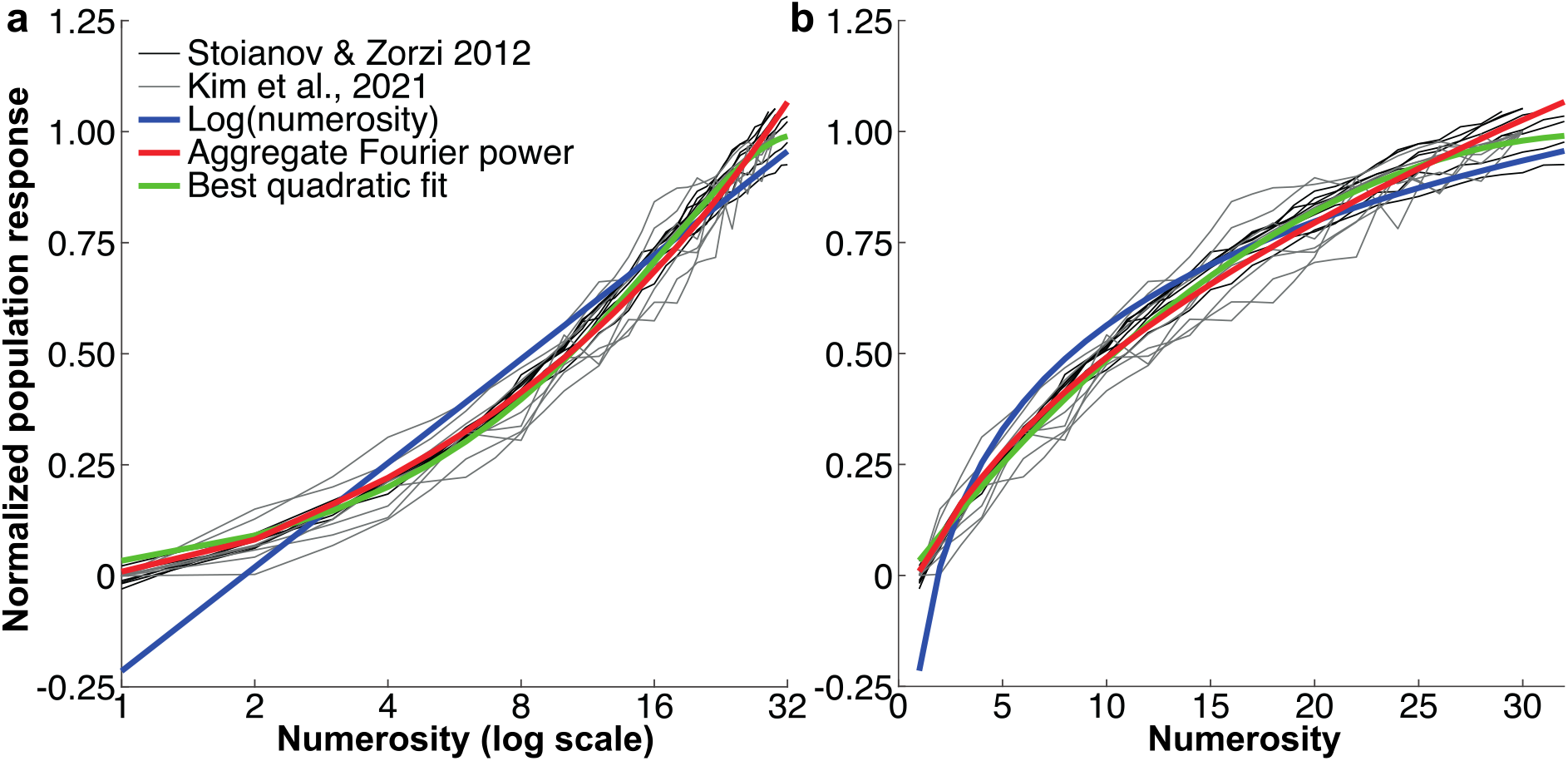
Comparison between the monotonic responses in neural network models of numerosity processing and the relationship of numerosity to aggregate Fourier power. **a** Shown on a log numerosity scale, following Stoianov and Zorzi (22). **b** Shown on a linear scale following Kim and colleagues (24). Responses shown in neural network studies (black and gray lines) are fit very closely by the relationship between numerosity and aggregate Fourier power (red), more closely than by log(numerosity) and similarly to the best quadratic fit to these responses.

### Differences in model fits within and between visual field maps

We separated each visual field map into two eccentricity ranges: near to the stimulus/fixation (below 1° eccentricity) and far from the stimulus/fixation (2–5.5° eccentricity). A linear mixed-effects model (fixed effects: visual field map, eccentricity range; random effects: participant) revealed that monotonically increasing Fourier power response models fit better near fixation (*p* = 4.4 × 10^-7^, *F*(1, 148) = 27.94), and differed between visual field maps (*p* < 10^-10^, *F*(16, 148) = 10.44). Post-hoc multiple comparisons revealed better model fits in the early visual field maps (V1, V2, V3, LO1) than elsewhere (Fig. 2c).

Numerosity tuned response model fits also differed between visual field maps (*p* < 10^- 10^, *F*(16, 133) = 10.79) in a similar linear mixed-effects model, but here post-hoc multiple comparisons showed poorer model fits in the early visual field maps than elsewhere. Also, unlike monotonically increasing Fourier power responses of early visual field maps, numerosity tuned response model fits did not differ significantly between near and far eccentricities (*p* = 0.195, *F*(1, 133) = 1.70). Therefore, progressions of tuned model fits with eccentricity were captured better by quadratic than sigmoid functions (Supplementary Fig. 5c).

### Monotonically decreasing responses outside early visual cortex

Some monotonically decreasing responses were seen outside early visual cortex. Average model fits across whole visual field maps did not differ significantly between monotonically decreasing and tuned numerosity models (one-way ANOVA, *p* = 0.281, *F*(1, 283) = 1.17). However, tuned models fit better (*p* = 0.0003, *F*(1, 49) = 14.86) within the previously described numerosity maps. Monotonically decreasing models generally fit better just outside the numerosity maps (Fig. 2a). Such responses are predicted by computational models of numerosity-tuned response derivation (21, 24), but typically at early stages preceding numerosity tuned responses, not alongside them (though see (23)). Alternatively, numerosity-tuned neural populations with preferred numerosities below one (38) would also decrease their responses as numerosity increases. These populations would be expected in the continuous topographic representation of numerosity, near populations with low numerosity preferences. We calculated the cortical distance between each recording site best fit by a numerosity tuned response and the nearest recording site best fit by a monotonically site’s preferred numerosity (Spearman rank order correlation, *ρ* = 0.585, *p* < 0.001; see Supplementary Fig. 11), suggesting monotonically decreasing sites are tuned, but with preferred numerosities below one.

## Discussion

We found monotonic increases in neural population response amplitude with increasing numerosity in the retinotopic locations of our stimuli, beginning in V1. While these monotonic responses follow numerosity closely, they are better predicted by aggregate Fourier power, which follows numerosity closely over a wide range of stimulus parameters for a fixed contrast. Monotonic responses shown in neural network studies of numerosity were also better predicted by aggregate Fourier power than by numerosity. Conversely, tuned responses overlapping with visual field maps in association cortices were not limited to the stimulus’s retinotopic location, and were better predicted by tuning for numerosity than aggregate Fourier power. We also found monotonically decreasing responses to numerosity near recording sites tuned to low numerosities, likely reflecting tuning for numerosities below one.

Numerosity is generally seen as a high-level visual feature or cognitive property, while Fourier power is a low-level representation of image contrast. Contrast energy at specific orientations and spatial frequencies drives V1 neurons’ responses (27). Therefore, the cortical response to any visual image begins with an approximate Fourier decomposition (27, 32). This transforms the visual image from the spatial domain (the image’s projection onto the retina) to the spatial frequency domain (a neural representation of the spatial frequency, orientation and phase of contrast). In the spatial domain, it does not seem possible to estimate numerosity regardless of item size and spacing: the area that must be integrated and the luminance, contrast or edge extent within that area change with item size and spacing.

However, numerosity is not directly estimated from the visual image, it is estimated from the early visual image representation, which is in the spatial frequency domain. Many functions in the spatial domain correspond to other functions in the frequency domain, and numerosity (spatial domain) corresponds to a nonlinear function of aggregate Fourier power (frequency domain) at a fixed contrast. This makes it potentially straightforward to estimate numerosity from the early visual, frequency domain image representation. Functional neuroimaging measures aggregate responses of large neural populations with a broad range of orientation and spatial frequency preferences (39). Aggregate Fourier power similarly sums contrast energy across orientations and spatial frequencies, and we propose this is why early visual neuroimaging responses follow numerosity with little effect of size or spacing: these early visual responses reflect aggregate Fourier power.

It is possible to generate phase-scrambled images that contain the same Fourier power distributions but with the locations of image contrast (an orthogonal phase component in the frequency domain) randomized. Such images yield strong responses in primary visual cortex, but even V2 responds poorly to such images as they lack the phase (position) structure found in natural images (40). Hard edges contain contrast at many frequencies with linked phases. Analysis of this phase structure may be important for object individuation. Therefore, we expect phase (position) structure to be required for numerosity-tuned responses to be derived from early visual, frequency domain image representations. As such, derivation of numerosity from frequency domain image representations simplifies object normalization processes that are required to disregard size and spacing, but is still compatible with object individuation processes and may also simplify these.

We sum Fourier power only within the first harmonic. This is a clear local minimum in the Fourier spectrum, but its spatial frequency varies with item size. The visual system seems unlikely to flexibly identify this limit and aggregate responses within it as we do in our analyses. We use this limit because the aggregate Fourier power in every harmonic (and so the total Fourier power to infinite spatial frequency) is proportional to the power in the first harmonic. For example, we show that the second harmonic’s power also follows numerosity closely, at half the amplitude. However, using a discrete Fourier decomposition on a pixelated image we cannot evaluate the power in further harmonic because they exceed the Nyquist frequency (a mathematical limit on which frequencies can be evaluated at a given sampling density). We therefore use a metric we can quantify that is equivalent to total Fourier power. The human visual system transforms images into the spatial frequency domain, but it does not use a discrete Fourier decomposition and its input is not pixelated: these are limitations imposed by computer models. Nevertheless, the visual system is still likely to have some spatial frequency limit. The finding that the aggregate response of early visual cortex is not affected by item size (25, 26) suggests it does not follow the Fourier power over a fixed spatial frequency range, but is proportional to aggregate Fourier power in the first harmonic and so to total Fourier power. It is unclear why this is so. It may be that the visual system samples the image very densely (certainly far more densely than 768 × 768 pixels as we do here) and little power falls beyond the frequencies it can evaluate. Alternatively, neurons responding to a specific spatial frequency and its harmonics will typically be activated together and are likely to interact. Although the nature of these interactions is unclear, responses to higher harmonics may be suppressed, for example. Finally, there are certainly differences between the visual system’s spatial frequency representation and a discrete Fourier decomposition, which is only a mathematical model. But it is clear that numerosity could be straightforwardly estimated from V1’s population response, as it is from a Fourier transform. The properties of lab numerosity displays in a Fourier transform and the Fourier decomposition’s close relationship to early visual spatial frequency analysis give an insight into how this computation could be achieved.

Aggregate Fourier power within the first harmonic follows numerosity closely because it is approximately constant with item size and spacing. Why is this constant? Increasing item size moves Fourier power to a narrower range of lower spatial frequencies (Supplementary Fig. 8). Its extent in the Fourier spectrum (bandwidth) follows 1/period, because spatial frequency is the inverse of spatial period. Conversely, Fourier power spectral density is high at these low frequencies because power spectral density of hard edges follows 1/frequency. Aggregate Fourier power, reflecting the product of bandwidth (1/period, i.e. frequency) and amplitude (1/frequency), is therefore constant.

Previous computational studies using neural networks have shown monotonically increasing responses to numerosity in a network that first decomposes the image using spatial receptive fields with surround suppression (22), as in the early visual system. This is conceptually similar to the transformation into the spatial frequency domain image representation that we describe. These networks’ monotonic responses are also spatially selective (22) and almost independent of item size (19), like we see here, when trained as generative models of numerosity displays. Even when network connections are randomly weighted, monotonic and numerosity-tuned units are found, suggesting that numerosity is reflected in image statistics (19, 24). Aggregate Fourier power may provide an effective statistic given that it is so straightforward to determine in spatial frequency domain image representations, and that the monotonic responses in these networks follow aggregate Fourier power more closely than they follow numerosity. Training to efficiently encode (but not discriminate) numerosity displays increases the discriminability of different numerosities from the network responses and reduces the influence of non-numerical properties of test displays (19). When transformed into the spatial frequency domain, numerosity displays (like the natural images biological visual systems learn from (41)) have the same 1/frequency power distribution that we propose underlies the size invariance of monotonic responses.

Therefore, this training may make network responses more sensitive to numerosity by incorporating this distribution into the spatial structure of network connections.

The aggregate Fourier power of a group of items is affected only slightly by its spacing. In Fourier decompositions, position becomes a phase component that we do not analyze. Like item size, spacing does not affect Fourier power because increasing the distance between items moves power to lower frequencies, with lower bandwidths and higher amplitudes. Nevertheless, the ratio of item size and spacing affects which frequencies fall within the first harmonic (Supplementary Fig. 9). If the distance between items is smaller than the item size, the between-item component of the Fourier spectrum falls outside the first harmonic, reducing power within this harmonic. Studies of numerosity perception avoid such crowding. If item spacing is very low (almost touching), the local minimum used to identify the first harmonic reflects group size rather than item size. Aggregate power then reflects one item (the group) rather than the group’s numerosity. Neither limitation would affect Fourier power if aggregated by the visual system over all frequencies.

On the other hand, item shape affects aggregate Fourier power considerably. Triangles have a greater aggregate Fourier power than other polygons (which have similar power to circles). Notably, their Fourier spectrum lacks a clear local minimum because a triangle’s sides are so far from parallel (Supplementary Fig. 10). Our procedure may therefore overestimate the first harmonic’s extent. Conversely, most stars have less aggregate Fourier power than polygons and circles. Stars contain higher frequency features that fall beyond the first harmonic. Again, neither limitation would affect Fourier power if aggregated by the visual system over all frequencies.

Numerosity estimation from aggregate Fourier power may explain several known effects of stimulus properties on numerosity perception. First, increasing item spacing slightly increases Fourier power and increases perceived numerosity (16). Similarly, tightly crowding items together reduces Fourier power and reduces perceived numerosity (34, 42, 43). Second, blurring items decreases their range of spatial frequencies, and reduces perceived numerosity (34). Third, complex shapes disrupt the relationship between numerosity and Fourier power. Such shapes affect both perceived numerosity (35) and early visual response amplitudes (36). So perceived numerosity, like V1 activation, depends on image properties.

Unlike Fourier decompositions, biological visual systems process different image locations with distinct neural populations. Our stimulus fell entirely in the central visual field. Humans can only integrate a limited spatial extent to estimate numerosity without making eye movements (44, 45). Very high numerosity stimuli must either be too large to see at a glance, be so dense that items are crowded, or use items too small to resolve. Aggregate Fourier power is unaffected by such limitations, so follows numerosity closely until at least 175. Human vision perceives such high numerosities differently to lower numerosities (46). So, the human visual system may approximate a Fourier decomposition to transform the image into the frequency domain, but has its own limitations.

Previous studies of non-numerical features in numerosity stimuli have focused on total item perimeter, area, density, or pattern extent. These follow numerosity in some stimuli (17, 18), but any single feature can be kept constant across numerosities. Numerosity estimation in the spatial frequency domain moves beyond this approach because item size and spacing have little effect here. Nevertheless, complex shapes, crowding, blurring, phase scrambling and contrast variations disrupt the relationship between numerosity and aggregate Fourier power. These factors therefore allow strong tests that a human or animal is responding to numerosity rather than aggregate Fourier power alone, which would otherwise give the appearance of numerosity-guided behavior and may itself be beneficial.

Some researchers have seen texture or spatial frequency metrics as non-numerical image features that can be used to perform numerosity discrimination tasks without true numerosity perception (13, 14, 47). Numerosity’s potentially straightforward estimation from the early visual spatial frequency domain image representation does not imply that humans perceive aggregate Fourier power rather than numerosity, but instead shows how numerosity itself could be straightforwardly estimated in the brain.

However, further processes are certainly involved in numerosity perception. There are situations where numerosity perception differs from true image numerosity. First, connecting items with lines or illusory contours (35, 48, 49) reduces perceived numerosity. This is generally thought to reflect higher-level grouping processes rather than image features (47). We show that lines or illusory contour inducers on items affect Fourier power, but this effect does not depend on whether a connection between items is formed. Conversely, in the connectedness illusion the reduction in perceived numerosity requires a connection. This is consistent with early visual EEG event related potentials, which initially reflect numerosity with no effect of connectedness, and are only affected by connectedness later (36). So, the connectedness illusion does not affect numerosity perception at the stage of numerosity estimation and is likely to reflect later processes. Similarly, numerosity adaptation affects numerosity perception (50) without affecting image content. This affects numerosity-tuned neural responses (51) but it is unclear whether early visual responses are also affected. Adaptation to the rate of finger tapping also affects visual numerosity perception (52) , and this effect seems very unlikely to arise in early visual cortex. Higher-level effects and contextual effects on numerosity perception are expected considering the extensive network of numerosity-tuned responses in human association cortices, which successively transform the representation of numerosity and include areas involved in attention, multisensory integration and action planning (5, 53).

Our model for estimating numerosity from aggregate Fourier power is conceptually similar to a spatial frequency analysis that Dakin and colleagues proposed to explain why numerosity perception is affected by density in very dense displays (14). That analysis uses the responses of high spatial frequency Laplacian Gaussian filters as a metric for numerosity. Such filters are not orientation-selective and do not transform the image into the spatial frequency domain, so do not closely model the response selectivity of neurons early visual cortex. Dakin and colleagues’ analysis was limited to items of a single size. Any such high spatial frequency filter’s response is strongly affected by item size (Supplementary Fig. 12a–c) while early visual cortex responses (25, 26), numerosity tuned responses (2, 4, 17) and numerosity perception (15, 16) are not. Therefore, this filter response predicts early visual responses poorly (Fig. 4a), as we have previously shown for numerosity-tuned responses (17, 18). Dakin and colleagues proposed that subjects perceive the response ratio of high and low spatial frequency filters, which is affected by density, rather than perceiving numerosity. Aggregate Fourier power is similarly affected by density. Their response ratio metric does not closely follow numerosity (Supplementary Fig. 12d) and also predicts early visual responses poorly (Fig. 4a), while aggregate Fourier power potentially allows straightforward estimation of numerosity itself. These differences are vital because humans rapidly and spontaneously perceive numerosity (15, 16). Therefore, while we were inspired by their insight that numerosity must be estimated from early visual responses, both of Dakin and colleagues’ proposed metrics for numerical vision predict neural responses and perception of numerosity poorly, particularly with respect to item size changes.

Models for subsequently computing numerosity-tuned responses rely on comparing monotonically increasing and decreasing responses (20, 21, 24), with their relative weights determining numerosity preferences. Human early visual population receptive fields have inhibitory surrounds (54), so populations with receptive fields further from stimulus area should monotonically decrease their responses with increasing numerosity. However, we observe no early visual monotonically decreasing responses, perhaps because negative fMRI responses have low amplitudes or fewer neurons with monotonically decreasing responses are needed (21). Using differently weighted excitatory and inhibitory synapses, tuned responses could be computed with monotonically increasing inputs only (20, 24). So, early visual neural populations alone may provide sufficient inputs to derive numerosity-tuned responses.

Numerosity-tuned responses emerge in neural network computational models trained to efficiently encode numerosity displays, or even if all weights are random (19, 24). If monotonic responses here arise from relationships between numerosity and image statistics (as we propose above), random weights from two resulting monotonic units could produce numerosity-tuned units with various numerosity preferences. These neural network models produce monotonic responses very early, by their second (22) or third (24) layers. Numerosity-tuned units can occur in the same layer (23), though in a feedforward deep convolutional network (24) numerosity-tuned units occur later, in the fourth layer, and are derived from responses of monotonic units in the third layer. Another deep convolutional neural network trained for object recognition (55) shows monotonic and numerosity-tuned units far later, in layers 11 and 13 respectively. Responses of earlier layers were not examined there, perhaps because only later units have inputs converging from the entire image. We kept our numerosity patterns small so that eye movements were not required to see the whole pattern clearly, and these patterns should easily fall within V1 population receptive fields. Late-stage responses in a ventral stream model do not seem to be a close model for either the monotonic responses of the human brain’s early visual cortex (25, 26) or the emergence of numerosity-tuned neurons in the lateral occipital cortex and their spread through the dorsal visual stream areas of the superior parietal lobule (5, 56). Nevertheless, both response types seem so straightforward to compute that they may also emerge in ventral stream areas to support object recognition processes. They may also emerge far later than early visual cortex if the items are spread over a large area (too big for early visual receptive fields), though even here the response of an early visual neural population should follow numerosity if averaged over many displays.

The transformation from monotonic to tuned responses that our results suggest also seems to transform Fourier power to numerosity. Which further processes would be needed to transform the early visual aggregate Fourier power response into a representation of numerosity that follows perceptual properties? We did not manipulate image contrast (i.e. dot darkness) in our fMRI stimuli, but Fourier power linearly decreases as contrast is reduced. It is well established that early visual neural response amplitudes depend on image contrast (57), so we expect that reducing image contrast would reduce V1 response amplitudes. Therefore, early visual responses are unlikely to follow numerosity (as others have proposed) as numerosity does not depend on contrast, and more likely to follow aggregate Fourier power (as we propose) as this does depend on contrast. Transforming early visual Fourier power responses into numerosity tuned responses would also require normalization for image contrast. Unlike early visual field maps, the responses of the first areas where we find numerosity tuned responses (visual field maps TO1 and TO2, i.e. area MT) are minimally effected by contrast (57). Therefore, contrast normalization at this stage would be sufficient to yield contrast-independent numerosity tuned responses. Global image contrast changes do not affect perceived numerosity until items become less visible (33). But contrast variations within a display may disrupt global normalization processes, potentially underlying the lower numerosity discrimination performance in contrast-varying displays compared to fixed contrast displays (34). Indeed, our results show that the range of aggregate Fourier power levels in different displays increased considerably as the range of contrasts increased, regardless of whether Fourier power is normalized by mean item contrast. The mean aggregate Fourier power across displays increased only slightly here, so any bias in perceived numerosity would likely be hard to detect given the large variation.

After contrast normalization, converting from aggregate Fourier power to numerosity may be as simple as including an exponential nonlinearity to compensate for sub-additive accumulation of Fourier power with numerosity. Nonlinear interactions between excitatory and inhibitory inputs to numerosity-tuned populations seem sufficient to implement this. This nonlinearity is the main difference between predictions of monotonically increasing responses to Fourier power and numerosity in our stimulus set. As a compressive nonlinearity, this might reflect fMRI response amplitude saturation with increasing neural activity. We believe this interpretation is unlikely because our numerosity stimuli produce response amplitudes far below the early visual cortex’s maximum fMRI response, and the same compressive nonlinearity is seen in monotonic responses of neural network models. We would not expect numerosity-tuned responses to show such saturation because they don’t increase response amplitude with numerosity.

Our experimental design cannot conclusively demonstrate that numerosity-tuned responses are derived from early visual frequency domain image representations, because we do not disrupt the early visual image representation and show effects on numerosity-tuned responses. Nevertheless, several findings suggest numerosity-tuned responses are computed from these early visual monotonic responses. First, almost all visual inputs to the cortex come through the primary visual cortex, which represents image features in the frequency domain and shows monotonic responses. There is no other known pathway through which numerosity-tuned neurons could be activated by visual inputs. Second, proceeding through the visual hierarchy, monotonic response model fits decrease in the same lateral occipital visual field maps where numerosity-tuned response model fits begin increasing, suggesting a transformation from monotonic to tuned responses. Third, computational models for numerosity-tuned responses (20–24) generally derive these responses from monotonic responses to numerosity. Our results differ from this process only by showing that early visual monotonic responses more closely follow frequency domain image properties, which also predict monotonic neural network responses more closely than numerosity does.

Unlike early visual monotonically increasing responses, numerosity tuned responses are not limited to neural populations with spatial receptive fields including the stimulus area. At the macroscopic scale, topographic maps of visual space and numerosity largely overlap, perhaps unsurprising as both are visually driven. This goes against the simplistic view of single brain areas having single functions. Indeed numerosity selectivity is found in many brain areas with diverse functions (5), many with spatial aspects: motion perception, spatial attention and eye movements (58–61). Numerical representations here may facilitate motion tracking, dividing attention and planning eye movements across multiple items respectively (5, 44, 45, 62, 63). But at a finer scale these response preferences are independent (5, 28), perhaps allowing neural responses to numerosity regardless of stimulus position (30). Linking specific numerosities and visual field positions would restrict all of these processes, and there is no link between particular numerosities and visual field positions in our stimuli or natural scenes. Conversely, to begin estimating numerosity from image contrast requires analysis of spatial responses in the stimulus area.

This distinction between the spatial selectivity of early visual monotonically responding populations and tuned populations in association cortices may reflect fundamental differences between the processes that estimate and use numerosity. Simpler animals like bees, zebrafish and newborn chickens display numerosity guided behaviors, but lack the complex numerical cognition of humans. Our findings suggest that apparently numerosity- guided behaviors in these animals may require only selective responses to orientation and spatial frequency in their simpler visual systems (64–66), with or without extracting numerosity from these spatial representations (67). The human brain appears to have built complex cognitive functions with inputs from these simple numerical responses (62).

## Methods

### Participants

We acquired fMRI data from eleven participants (aged 25–39 years, one female, one left-handed). All had normal or corrected-to-normal visual acuity, good mathematical abilities and were well educated. Participants were familiarized with the numerosity stimuli using tasks that required numerosity judgments before scanning. Written informed consent was obtained from all participants. All experimental procedures were approved by the ethics committee of University Medical Center Utrecht.

Data from participants P1-P5 were included in a previous study (5), although we use updated preprocessing protocols here. These participants were scanned while viewing all four stimulus configurations (Fig. 1), to test for responses to non-numerical features over a broad range of stimulus parameters. P6-P11 were only scanned while viewing the constant area and constant size configurations: previous studies show very similar responses across stimulus configurations, and non-numerical features that differ between configurations predict numerosity selective (17) and monotonic (25) responses poorly.

### Visual stimuli

Following experimental protocols described previously (4, 5, 28), we acquired 7T fMRI data while participants viewed numerosity and visual field mapping stimuli. All stimuli were presented by back-projection on a 15 × 9 cm screen inside the MRI bore, which participants viewed through prism glasses and a mirror attached to the head coil. Distance from the participants’ eyes to the display was 41 cm, with a visible resolution of 1,024 × 538 pixels.

Visual stimuli were generated using MATLAB and Psychtoolbox-3 (68). Stimuli were sets of black circles (items) on a middle gray background. A diagonal cross of thin red lines intersected the center of the screen and covered the entire display throughout the experiment to aid with accurate fixation.

Numerosity stimuli consisted of groups of black dots (items) within 0.75° (visual angle, radius) of a fixation cross at the center of the display. We used four stimulus configurations with different progressions of item size and spacing with numerosity (Fig. 1). The first three kept total item area (and display luminance), individual item size, or total item perimeter constant. These placed items randomly but approximately homogeneously within the stimulus area. The fourth (“high density”) grouped the items constant total item area entirely inside a 0.375° radius circular area, randomly placed inside the stimulus area. 10% of displays showed white items instead of black, and participants responded to these displays with a button press (80-100% accurately in each run). No numerosity judgments were required.

The numerosities one through seven were first presented in ascending order, with numerosity changing every 4200 ms (two TRs, see Supplementary Fig. 1c–d inset). Within this period, a numerosity pattern was shown for 300 ms, alternating with 400 ms of gray background, repeated six times. Each numerosity pattern had items drawn in new, random positions. These short presentations prevented participants from counting. Following this, twenty items were presented in the same way for eight TRs (16.8s, 24 presentations of a pattern). These periods of twenty items served to distinguish between very small and very large tuning widths (4, 69). Then numerosities one through seven were presented as before, but in descending order, followed by another long period of twenty items. This cycle was repeated 4 times in each scanning run.

Repeated presentation of stimuli with the same numerosity likely caused some adaptation (50) or repetition suppression. To minimize effects of adaptation on estimated response models, we used a single model to summarize responses to both increasing and decreasing numerosity. This ensured each numerosity presentation was preceded by stimuli that caused both higher and lower responses, reducing dependence on preceding stimuli.

In a separate scanning session, visual field mapping was used to delineate visual field maps and determine the position selectivity of our recording sites, following protocols described previously (69, 70). Briefly, a bar filled with a moving checkerboard pattern stepped across a 6.35° (radius) circle in the display center in eight (cardinal and diagonal) directions. Participants fixated the same central fixation cross, pressing a button when this changed color to ensure fixation and attention.

### fMRI acquisition and data pre-processing

We acquired MRI data on a 7T Philips Achieva scanner, following protocols described fully in our previous studies (5, 28, 71). Briefly, we acquired T1-weighted anatomical scans, automatically segmented these with Freesurfer, then manually edited labels to minimize segmentation errors using ITK-SNAP. This provided a highly accurate cortical surface model at the gray-white matter border to characterize cortical organization. We acquired T2*- weighted functional images using a 32-channel head coil at a resolution of 1.77 × 1.77 × 1.75 mm, with 41 interleaved slices of 128 × 128 voxels. The resulting field of view was 227 × 227 × 72 mm. TR was 2100 ms, TE was 25 ms, and flip angle was 70 degrees. We used a single shot gradient echo sequence with SENSE acceleration factor 3.0 and anterior-posterior encoding. This covered most of the brain but omitted anterior frontal and temporal lobes, where 7T fMRI has low response amplitudes and large spatial distortions. Each functional run acquired 182 images (382.2s) of which the first six (12.6s) were discarded to ensure a steady signal state. In each scan session, we acquired six to eight repeated runs in one stimulus configuration, plus a top-up scan with the opposite phase-encoding direction to correct for image distortion in the gradient encoding direction (72), and a T1-weighted anatomical image with the same resolution, position and orientation as the functional data. Different stimulus configurations were tested in different sessions.

Co-registration of functional data to the high-resolution anatomical space was performed using AFNI (afni.nimh.nih.gov; (73)), which differs from our previous studies. A single transformation matrix was constructed, incorporating all the steps from the raw data to the cortical surface model to reduce the number of interpolation steps to one. No other spatial or temporal smoothing procedures were applied. A T1 image with the same resolution, position and orientation as the functional data was first used to determine the transformation to a higher resolution (1 mm isotropic) whole-brain T1 image (3dUnifize, 3dAllineate). For the fMRI data, we first applied motion correction to two series of images that were acquired using opposing phase-encoding directions (3dvolreg). Subsequently, we determined the distortion transformation between the average images of these two series (3dQwarp). We then determined the transformation in brain position between and within functional scans (3dNwarpApply). Then we determined the transformation that co-registers this functional data to the T1 acquired in the same space (3dvolreg). We applied the product of all these transformations to every functional volume to transform our functional data to the whole-brain T1 anatomy. We repeated this for each fMRI session to transform all their data to the same anatomical space.

We then imported these data into mrVista (github.com/vistalab/vistasoft). We averaged visual field mapping scan runs. We averaged scans in the same numerosity stimulus configuration together. For each stimulus configuration we also averaged data from odd runs and averaged data from even runs for cross-validation. Within these averaged runs, we averaged repeated stimulus cycles.

### Analyses of visual features co-varying with numerosity

Following previously described methods (17, 18), we quantified several non-numerical features of each display in each numerosity stimulus configuration using methods analogous to those described fully in (17). For monotonic models, we tested predictions of responses monotonically increasing or decreasing linearly or logarithmically (whichever fit best) with: [1] numerosity; individual item [2] area and [3] perimeter; total item [4] area (74) and [5] perimeter (75); [6] area and [7] perimeter of the convex hull (76); density of [8] luminance, [9] edges, and [10] number within the convex hull; root mean square contrast [11] across the display and [12] in the convex hull; and [13] aggregate Fourier power. We also tested a model proposed to capture numerosity perception using the responses of a high spatial frequency Laplacian of Gaussian center-surround filter (14). We used a filter with a 2- pixel standard deviation here, but other filter sizes made very similar predictions (Supplementary Fig. 12). We convolved this filter with each image and summed the response over all image locations. We also tested the ratio of responses of high and low spatial frequency filters (the response ratio model (14)). We used filters with standard deviations of 1 and 34 pixels here as the prediction of this filter pair was most strongly correlated with numerosity in our displays. We also tested spatial pRF models following the visual positions of [14] edges and [15] luminance, summing these across all displays of the same numerosity.

Additionally, for each display we determined the aggregate Fourier power within the first harmonic using MATLAB functions. We first normalized each image to a value of zero on the background and one in the items. We took a two-dimensional Fourier transform of the resulting image (fft2) and shifted the zero-frequency component to the center (fftshift) (Fig. 3a). We then took annular frequency bins (at all orientations) at intervals of 1 cycle/image (excluding zero frequency), in which we summed the absolute power spectral density (PSD) of the Fourier transform. Plotting PSD across frequencies (Fig. 3b) identifies a clear local minimum at the end of the first harmonic. We identified this by finding the lowest frequency above the frequency of the global maximum PSD where: either the first or second derivative of PSD reaches its maximum and the PSD is below 25% of the global maximum PSD. This effectively identifies the sharpest change in PSD, i.e. the end of the first harmonic, avoiding artefacts resulting from the image pixelation or Fourier transform discretization. We summed PSDs of all frequency bins until this frequency to give the aggregate Fourier power in the first harmonic for each display. We also determined how well the relationship between aggregate Fourier power and numerosity generalizes across a much greater range of item sizes, spacings and shapes that were not included in our fMRI experiments. Display resolution affects Fourier power. We arbitrarily evaluated our stimuli’s power in 768 × 768- pixel images but divided it by 768 × 768 × 2^0.5^ to generalize to any resolution. This makes aggregate Fourier power in the first harmonic very nearly one for one-item displays.

We used the following MATLAB code to determine the aggregate Fourier power in the first harmonic, from images placed in the image buffer using Psychtoolbox-3 (68):

~~~
ItemColor = 0; %Intensity values in the first color channel
BackgroundColor = 128;
GetImageSize = 768; %Image must be square, must be an even number of pixels

%Get image from image buffer
NumImage = Screen(’GetImage’, display.windowPtr, [display.numpixels(1)/2-
GetImageSize/2 display.numpixels(2)/2-GetImageSize/2
display.numpixels(1)/2+GetImageSize/2
display.numpixels(2)/2+GetImageSize/2]);

%Normalise image so background is zero and item is one
NumImage = (BackgroundColor-single(NumImage(:,:,1)))./(BackgroundColor - ItemColor);

%Take 2D Fourier transform, shift zero frequency component to centre
FourierImage = abs(single(fftshift(fft2(NumImage, GetImageSize, GetImageSize))));

%Sum over all orientations for each frequency
[x,y] = meshgrid(-GetImageSize/2:GetImageSize/2-1, -GetImageSize/2:
GetImageSize/2-1);
[∼,Radius] = cart2pol(x,y);
Radius = round(Radius);
FreqBins = zeros(1,(GetImageSize/2));
for RadCounter = 1:(GetImageSize/2)
    FreqBins(RadCounter) = sum(FourierImage(Radius==RadCounter));
end

%Find the limit of the first harmonic. For robustness, use the 1st and 2nd derivative, and use whichever returns the lowest frequency.
[maxVal, maxPos] = max(FreqBins);
deriv1 = [diff(FreqBins)];
deriv1(FreqBins>(maxVal/4)) = 0;
deriv1(1:maxPos) = 0;
[∼, whichMin1] = max(deriv1);

deriv2 = [0 diff(diff(FreqBins))];
deriv2(FreqBins>(maxVal/4)) = 0;
deriv2(1:maxPos) = 0;
[∼, whichMin2] = max(deriv2);
whichMin = min([whichMin1 whichMin2]);

%Sum to this limit to get the aggregate Fourier power in the first harmonic
AggregatePower = sum(FreqBins(1:whichMin));

%Compensate for the image resolution
AggregatePower = AggregatePower/(GetImageSize^2*sqrt(2));

%To convert this to a numerosity estimate if preferred. This is somewhat approximate and descriptive of the observed relationship.
NumerosityEstimate = AggregatePower^2.2;
~~~

We also determined how well the relationship between aggregate Fourier power and numerosity generalizes across a much greater range of item sizes, spacings and shapes that were not included in our fMRI experiments. We tested single circles from 1-240 pixels diameter placed at the image center. We then tested images of 7 circles, each of diameter 16 pixels, spaced randomly but evenly within a 50 to 528 pixel diameter circular area. The minimum center-to-center distance increased with the group area. We then tested images of 16-pixel diameter circles from numerosity 1-175 within a 700-pixel diameter circular area, placed at least 24 pixels apart (edge-to-edge).

We then tested regular convex polygons from 3 corners (equilateral triangle) to 10 corners (regular decagon), plus a circle (i.e. infinite corners), with all corners 30 pixels (radius) from the image center. We also tested regular stars with 3 to 10 points (at 30 pixels radius), interleaved with concave corners at 10 pixels radius. Spatial frequency differs with orientation in these shapes, so we determined the frequency of the first harmonic separately at each orientation and summed the PSD within the resulting area.

We finally tested pairs of dots with and without connecting elements to determine whether the effects of these connecting elements on Fourier power predict their effects on perception. All dots we 30 pixels in diameter. All bars were 4 pixels wide and the same length as the (edge-to-edge) distance between dots. Illusory contour inducers were 4 pixels wide, the same color as the background, and extending 10 pixels into the dot. To separate effects of connected configurations from changes to image properties, we included conditions where the bar or illusory contour inducer was split in the middle and rotated around the dot center.

To model predictions of the mean fMRI response to aggregate Fourier power (or any other non-numerical feature) we took the mean value over all displays of the same numerosity and stimulus configuration.

### Neural response models

For each stimulus configuration’s data and cross-validation splits, we fit several candidate response models testing how well a putative neural population with monotonic or tuned responses to each stimulus feature predicts the observed responses. For monotonic response models, we convolved the sequence of presented feature amplitudes with a canonical hemodynamic response function (HRF) to give a predicted fMRI time course. We found the optimal scaling for this prediction using a general linear model (Supplementary Fig. 1) to determine the baseline and *β* (scaling factor) for each recording site. This revealed whether responses increased or decreased with feature amplitude (the sign of *β*) and the response variance explained by the scaled prediction (R^2^). Log(numerosity) predicts early visual EEG (25, 31, 77) and fMRI (26) response amplitudes well. To give each non-numerical feature the best possible chance of explaining these responses, we tested both logarithmic and linear scaling and chose that which gave the highest variance explained across all recording sites and participants. For each non-numerical feature, the best performing model (logarithmic or linear) was consistent with that chosen to best predict numerosity tuned responses (17, 78).

We quantified these model fits under cross-validation between odd and even scanning runs per condition: we fit the model’s free parameters on one half of the data and evaluated the resulting model’s fit on the complementary half. This primarily compensates for the free parameters in the visual position selective pRF models and the tuned models: the monotonic models have no free parameters. Using cross-validated model fits allows an unbiased comparison between the monotonic model fits, visual position selective pRF model fits, and tuned model fits. We refit the scaling factor to evaluate the prediction, as scaling factors change between scans and sessions, but did not allow the scaling factor of monotonic models to change sign.

We then compared the cross-validated variance explained in visual field maps V1-V3 between the monotonic numerosity model and alternative monotonic and visual position selective pRF models. We first excluded from this comparison any recording site where no model explained more than 20% of the response variance (i.e. R^2^<0.2) before cross-validation That is, we selected voxels based on model fits in the data on which the model was fit, rather than that on which the model was evaluated, to avoid circularity between voxel selection and model comparison. We also avoid by selecting voxels where any compared model performs well, rather than voxels where one specific model performs well. We then took the median cross-validated fit from each hemisphere (across V1-V3), and from both cross-validation halves. To compare any two models, we then took the paired difference between these models’ fits in each hemisphere, resulting in a distribution of differences in model fits. We determined whether this distribution showed a significant difference between model fits using Wilcoxon signed-rank tests, as this population of fit differences was not normally distributed. We then corrected the resulting p-values for multiple comparisons using false discovery rate correction (37).

We calculated the proportion of variance in the responses to all stimulus configurations (Area, Size, Perimeter, Density) that these models explained, rather than simply averaging the variance explained values from each stimulus configuration. This ensured that configurations with larger fMRI signal variance were weighted more heavily in the average across stimulus configurations. Arbitrary differences in fMRI response amplitude between scan sessions (79) prevented us from quantifying differences in neural response amplitude between stimulus configurations.

We estimated tuned response models as previously described (4, 5, 17), following a pRF modeling approach (69). For tuned response models, we only compared predictions of tuning for numerosity and aggregate Fourier power: we have previously demonstrated that numerosity tuning predicts responses better than tuning for other features (5, 17). Briefly, for each recording site, we start with a large set of candidate neural response models describing numerosity or Fourier power tuning using logarithmic Gaussian functions (3–5) characterized by two parameters: (1) a preferred numerosity or Fourier power (mean of the Gaussian distribution) and (2) a tuning width (standard deviation of the Gaussian). Candidate neural response time courses reflected the amplitude of the Gaussian at the numerosity or Fourier power presented at each time point. We convolved these with an HRF to generate candidate fMRI time courses. We found the candidate fMRI time course best correlated with the response at each recording site, giving the neural response model parameters that best predict that recording site’s response and the response variance explained by that prediction (R^2^).

We restricted candidate preferred numerosities to be between 1.05–6.95, within the range of numerosities presented. Beyond the tested range, tuning function parameters cannot be determined accurately (4, 5, 69) and neural response amplitude predictions monotonically increase or decrease within the presented numerosity range, complicating comparisons to monotonic models.

We used the same procedure to compare the cross-validated variance explained in the numerosity maps between the numerosity-tuned model and aggregate Fourier power tuned model, as we have previously demonstrated that the numerosity-tuned model fits better than models tuned to other non-numerical visual features.

### Visual field mapping analysis

Visual field mapping data were analyzed following a standard pRF analysis (69, 70). We identified visual field map borders based on reversals in the cortical progression of the polar angle of recording sites’ visual field position preferences, drawing these by hand on the cortical surface of an inflated rendering of each participant’s brain (Fig. 2a and Supplementary Figs. 3 and 4). As well as the early visual field maps (V1, V2, V3), we identified higher visual field maps in the lateral/temporal occipital (LO1-LO2, TO1-TO2), parietal association (V3A/B, IPS0-IPS5) and frontal (sPCS1-sPCS2, iPCS) cortices with reference to landmarks identified in previous studies (59, 80–82).

We binned recording sites in each visual field map by pRF eccentricity, at 0.2° intervals from 0° to 5.5°. We took the mean and standard error of the variance explained by each response model among the recording sites within each bin in plotted this against bin eccentricity. We fit relationships between bin mean variance explained in each numerosity response model and eccentricity using cumulative Gaussian sigmoid functions with 4 parameters: the point of inflection (cumulative Gaussian mean), slope (standard deviation), maximum asymptote (variance explained at fixation) and minimum asymptote (variance explained at 5.5° eccentricity). We also fit quadratic curves to these, with three parameters: intercept, slope and quadratic term. This includes the possibility of a constant relationship, linear relationship, and more complex relationships that allow for an increase in variance explained in the middle of the eccentricity range, which was often observed. For both fit functions, we bootstrapped these fits using 1000 samples drawn from the unbinned variance explained data across all participants. We took the median of these bootstrapped fit parameters as the best fitting curve. We determined 95% confidence intervals from the 2.5 and 97.5 percentile from the distribution of bootstrapped fits at each eccentricity. For each visual field map and response model, we chose the function that gave the best correlation between the fit and the bin means.

### Analysis of relationship between spatial and numerical responses

We also analyzed how variance explained by the best fitting response models was related to recording sites’ spatial pRFs. We use pRF eccentricity (distance from the visual field center, where numerosity stimuli were presented) to quantify the pRF’s coverage of the stimulus area: pRF size linearly follows eccentricity (69, 70), so pRFs with shared eccentricities cover the numerosity stimulus area similarly. For each visual field map, we binned all participants’ recording sites by eccentricity. We fit relationships between bin mean variance explained and eccentricity using a sigmoid cumulative Gaussian function (i.e. variance explained decreasing with eccentricity) and a quadratic function (allowing many relationships between variance explained and eccentricity). For each visual field map and response model, we chose the function that best fit the bin means.

### Analysis of differences in model fits within and between visual field maps

We used linear mixed effects models to determine how the goodness of fit of our monotonic and tuned response models differed between visual field maps and eccentricity ranges. These models included visual field map and eccentricity range as fixed factors and participant as a random factor, because the quality of fMRI data varies between sessions and participants. Marginal tests for the fixed effects were adjusted using Satterthwaite degrees of freedom approximation (83). To determine which visual field maps showed different goodness of fit, we used post-hoc multiple comparisons as part of the linear mixed effects statistical model. These are corrected for multiple comparisons by using Tukey’s honestly significant difference test (84), which gives the marginal means and confidence intervals shown in Fig. 2c.

### Comparison to neural network models

To compare the predictions of monotonic responses to aggregate Fourier power against the responses of neural network models, we first extracted the data from Stoianov and Zorzi (22) (their Supplementary Figure 4A) and Kim and colleagues (24) (their Figure 5B) using graphreader.com. Each of these figures shows several lines, so we repeated this for 8 lines from each figure, evaluating these lines at each integer shown. Stoianov and Zorzi show responses on a log numerosity axis without normalization, while Kim and colleagues show responses on a linear numerosity axis normalized from zero (for the smallest response seen, to numerosity 1) to one (for the largest response seen, to numerosity 30). We normalized the responses shown by Stoianov and Zorzi globally following Kim’s approach (the mean response to numerosity 1 was normalized to 0, the mean response to numerosity 30 was normalized to 1) , though we kept the differences between different lines’ start and end points. We then plotted the data from both studies on the same axes with both log and linear scales. Finding that the monotonic responses shown by these studies are very similar, we subsequently treated them as one set of 16 lines. While we could not determine the aggregate Fourier power of the displays used in these studies, our own results show that item size and spacing have minimal effects on aggregate Fourier power. Therefore, we took the nonlinear function of numerosity predicted by aggregate Fourier power in our own displays and linearly scaled this to fit all the data points. We similarly scaled the log(numerosity) function. Finally, we fit a quadratic function (second-degree polynomial) to all these data. Then, for each of the 16 lines shown in the neural network studies we calculated the correlation coefficient to each of these 3 functions. We performed paired *t*-tests on the sets of 16 correlation coefficients from each function.

## Data availability

The data sets generated during the current study are available from the corresponding author upon reasonable request.

## Code availability

The code that supports the findings of this study is available from the Vistasoft repository (https://github.com/vistalab/vistasoft).

## Acknowledgments

This work was supported by the Netherlands Organization for Scientific Research (452.17.012 to B.M.H.); and the Portuguese Foundation for Science and Technology (IF/01405/2014 to B.M.H.). We thank Serge Dumoulin for sharing data collected by his lab.

## Author contributions

Conceptualization, J.M.P. and B.M.H.; Methodology, B.M.H.; Software, J.M.P., M.A., and B.M.H.; Validation, J.M.P. and B.M.H.; Formal Analysis, J.M.P. and B.M.H.; Investigation, J.M.P. and B.M.H.; Data Curation, J.M.P., M.A., T.C.C., and B.M.H.; Writing – Original Draft, J.M.P. and B.M.H.; Writing – Review & Editing, J.M.P., M.A., T.C.C., and B.M.H.; Visualization, J.M.P. and B.M.H.; Supervision, B.M.H.; Project Administration, B.M.H.; Funding Acquisition, B.M.H.

