## Supplementary Information for "Numerosity tuning in human association cortices and local image contrast representations in early visual cortex"

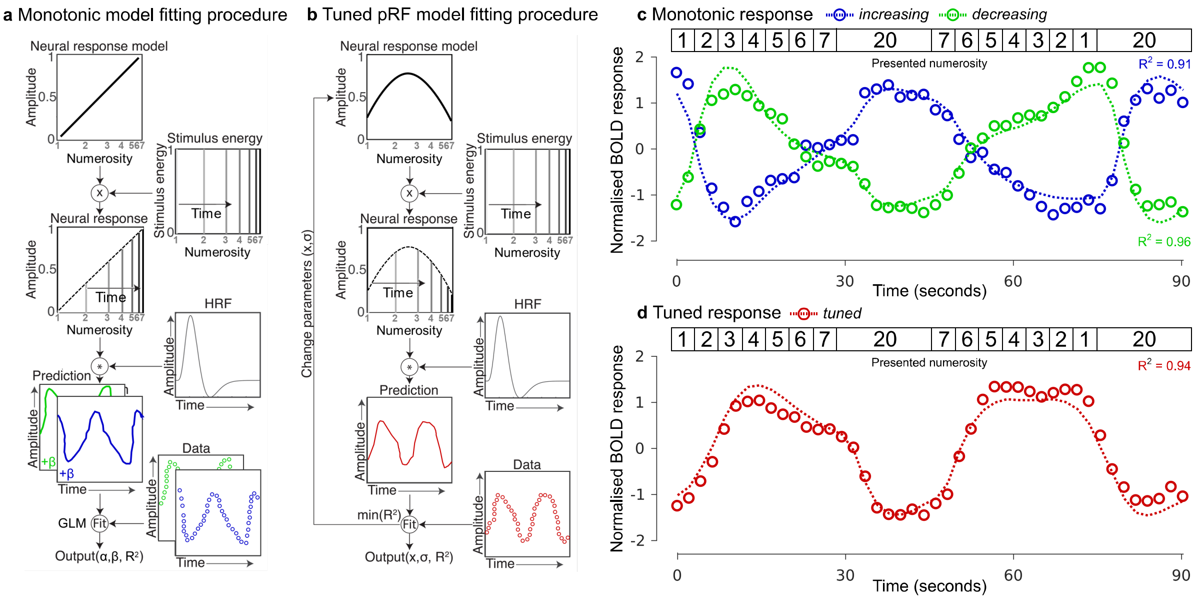


**Supplementary Fig. 1: Model fitting. a, b** Monotonic model and population receptive field (pRF) fitting procedures for estimating monotonic and tuned response functions for numerosity, respectively. **c, d** Illustrative neural response time course (circles) and model prediction (dashed line) for three separate voxels best described by preferences for monotonically increasing (blue), monotonically decreasing (green) or tuned (red) responses to numerosity. Similar models were used to test predictions of response models based on non-numerical stimulus features. Numerosity varied in ascending and descending order for all stimulus configurations to counterbalance adaptation effects (see stimulus inset).

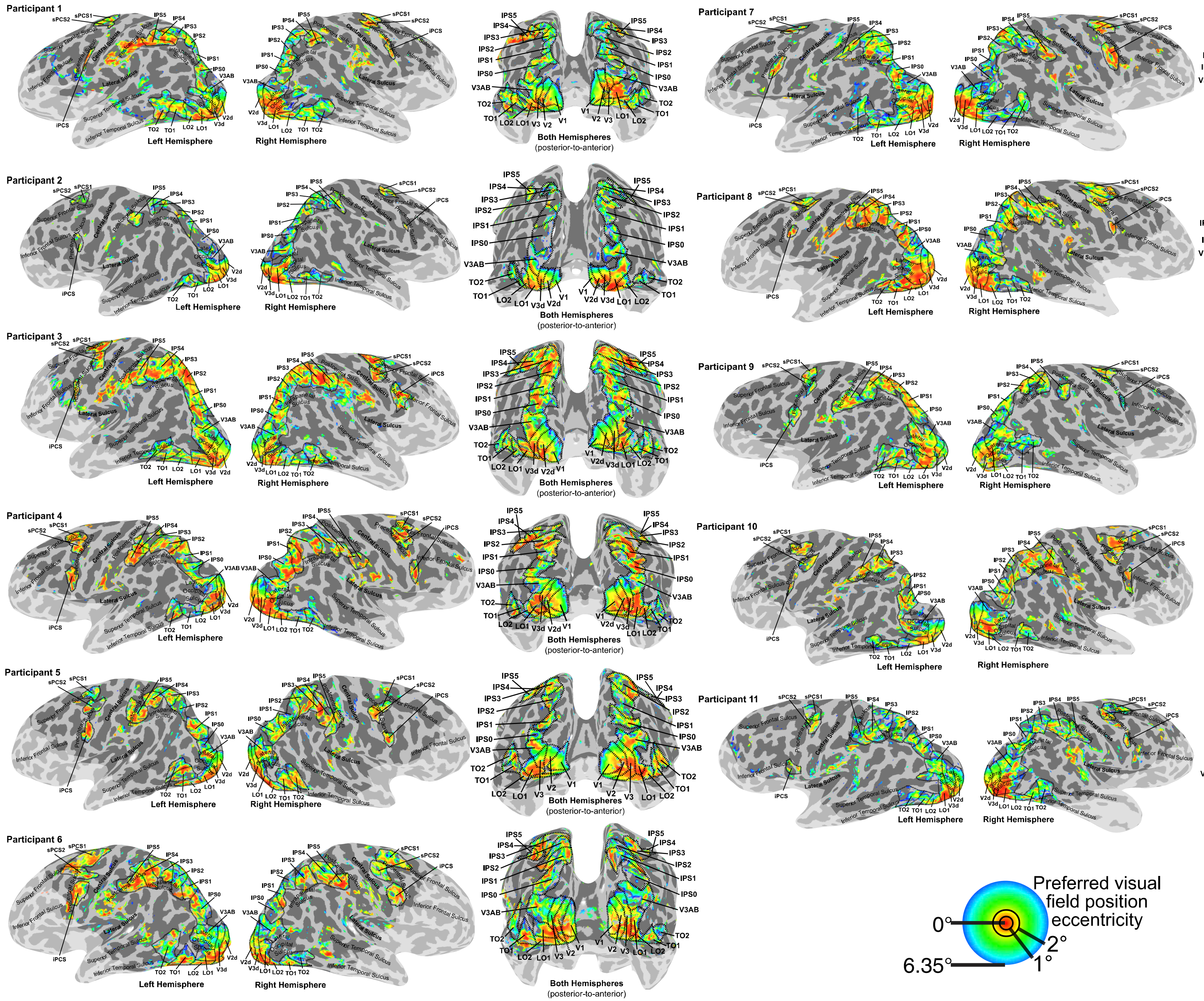

Participant 1
Left Hemisphere
Right Hemisphere
Both Hemispheres
(posterior-to-anterior)
Participant 2
Left Hemisphere
Right Hemisphere
Both Hemispheres
(posterior-to-anterior)
Participant 3
Left Hemisphere
Right Hemisphere
Both Hemispheres
(posterior-to-anterior)
Participant 4
Left Hemisphere
Both Hemispheres
(posterior-to-anterior)
Participant 5
Left Hemisphere
Right Hemisphere
Both Hemispheres
(posterior-to-anterior)
Participant 6
Left Hemisphere
Right Hemisphere
Both Hemispheres
(posterior-to-anterior)
Participant 7
Left Hemisphere
Right Hemisphere
Both Hemispheres
(posterior-to-anterior)
Participant 8
Left Hemisphere
Right Hemisphere
Both Hemispheres
(posterior-to-anterior)
Participant 9
Left Hemisphere
Right Hemisphere
Both Hemispheres
(posterior-to-anterior)
Participant 10
Left Hemisphere
Right Hemisphere
Both Hemispheres
(posterior-to-anterior)
Participant 11
Left Hemisphere
Right Hemisphere
Both Hemispheres
(posterior-to-anterior)
Preferred visual
field position
eccentricity
0°
2°
1°
6.35°

**Supplementary Fig. 2:** Preferred visual field position eccentricities determined from the visual field mapping task. All colored sites have greater than 10% model variance explained ($R^2$>0.1). Dashed black lines show visual field map borders and black labels highlight visual field map names. The light shaded region is outside the fMRI recording volume.

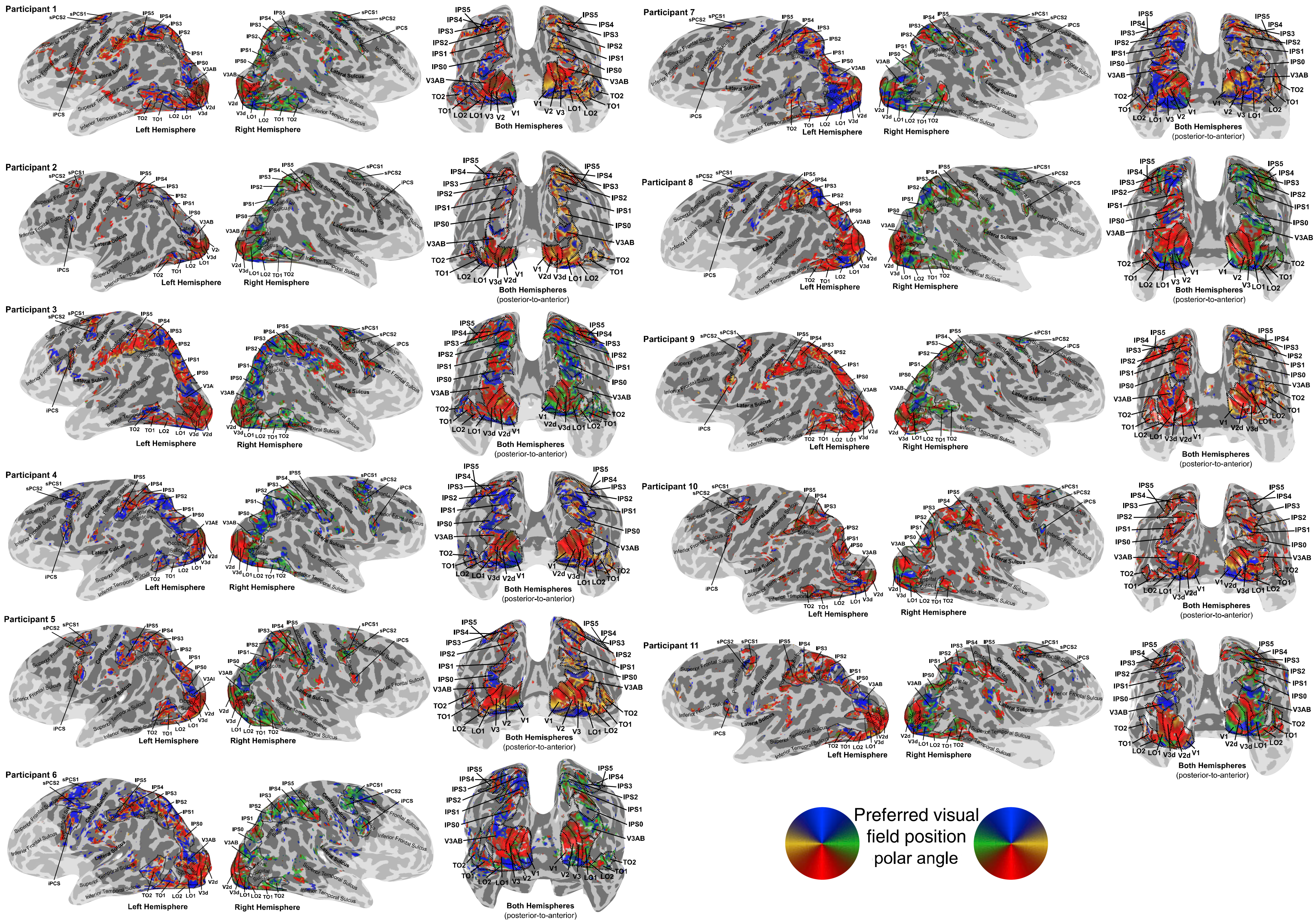

Participant 1
Left Hemisphere
Right Hemisphere
Both Hemispheres
Participant 2
Left Hemisphere
Right Hemisphere
Both Hemispheres
(posterior-to-anterior)
Participant 3
Left Hemisphere
Right Hemisphere
Both Hemispheres
(posterior-to-anterior)
Participant 4
Left Hemisphere
Right Hemisphere
Both Hemispheres
(posterior-to-anterior)
Participant 5
Left Hemisphere
Right Hemisphere
Both Hemispheres
(posterior-to-anterior)
Participant 6
Left Hemisphere
Right Hemisphere
Both Hemispheres
(posterior-to-anterior)
Participant 7
Left Hemisphere
Right Hemisphere
Both Hemispheres
(posterior-to-anterior)
Participant 8
Left Hemisphere
Right Hemisphere
Both Hemispheres
(posterior-to-anterior)
Participant 9
Left Hemisphere
Right Hemisphere
Both Hemispheres
(posterior-to-anterior)
Participant 10
Left Hemisphere
Right Hemisphere
Both Hemispheres
(posterior-to-anterior)
Participant 11
Left Hemisphere
Right Hemisphere
Both Hemispheres
(posterior-to-anterior)
IPS5
IPS4
IPS3
IPS2
IPS1
IPS0
V3AB
TO2
TO1
LO2
LO1
V3
V3d
V2
V2d
V1
sPCS1
sPCS2
iPCS
Central Sulcus
Superior Frontal Sulcus
Inferior Frontal Sulcus
Lateral Sulcus
Superior Temporal Sulcus
Inferior Temporal Sulcus
Preferred visual
field position
polar angle

**Supplementary Fig. 3:** Preferred visual field position polar angles determined from the visual field mapping task. All colored sites have greater than 10% model variance explained ($R^2$>0.1). Dashed black lines show visual field map borders and black labels highlight visual field map names. The light shaded region is outside the fMRI recording volume.

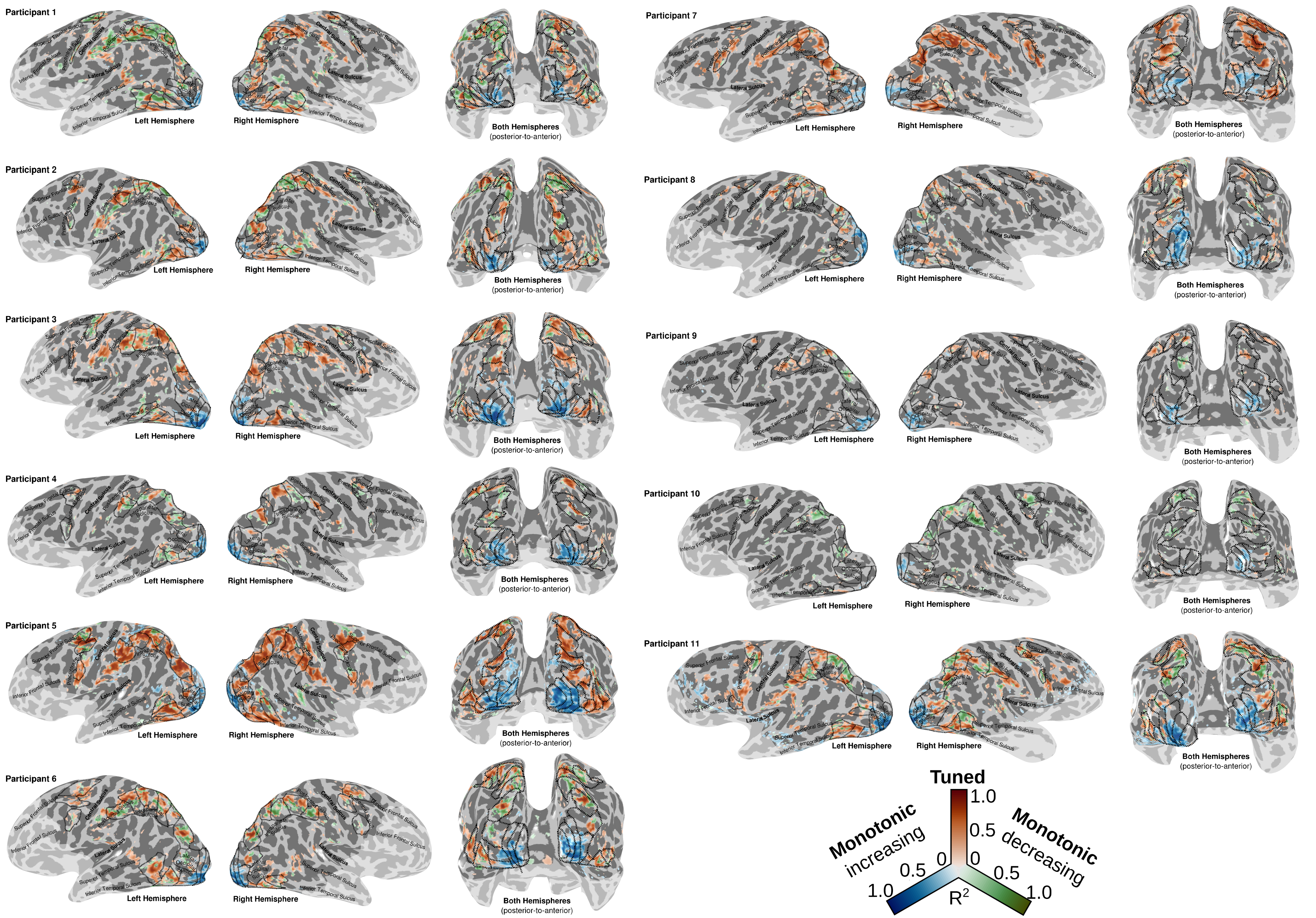
Participant 1
Left Hemisphere
Right Hemisphere
Both Hemispheres
(posterior-to-anterior)
Participant 2
Participant 3
Participant 4
Participant 5
Participant 6
Participant 7
Participant 8
Participant 9
Participant 10
Participant 11
Tuned
1.0
0.5
0
Monotonic
increasing
Monotonic
decreasing
R2

**Supplementary Fig. 4:** Distribution of best-fitting numerosity response model across the cortical surface for all subjects: winning response model is monotonically increasing (blue), monotonically decreasing (green), or tuned (red). Color intensity shows variance explained by winning response model. All colored sites have greater than 20% model variance explained ($R^2$>0.2). Data are cross-validated model fits from odd and even scanning runs. Dashed black lines show visual field map borders. The light shaded region is outside the fMRI recording volume.

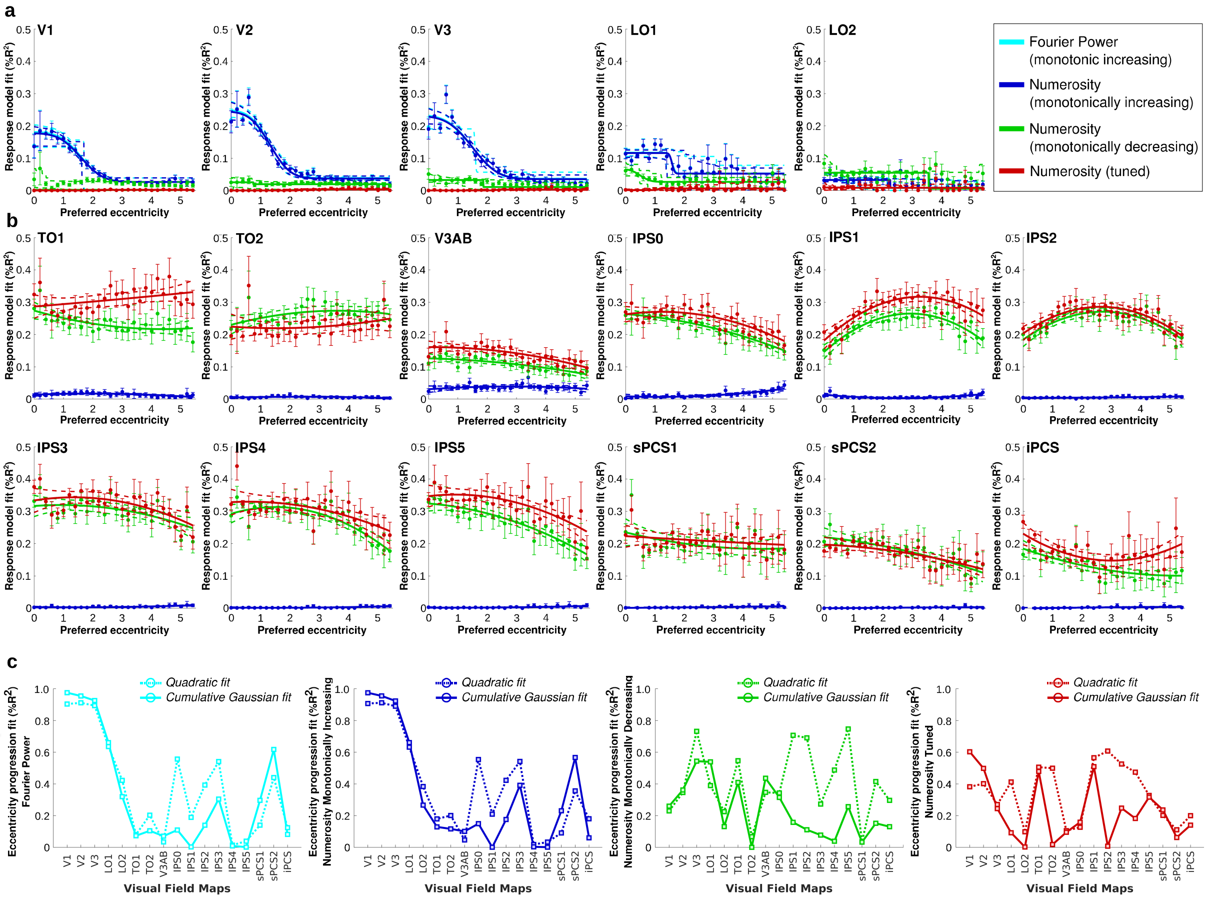


**Supplementary Fig. 5: Monotonically increasing responses in early visual cortex are location-dependent**. Progression of eccentricity preference plotted against different numerosity and Fourier power model fits across visual field maps. Data are cross-validated model fits from odd and even scanning runs. Filled circles show mean variance explained per eccentricity bin, error bars show the standard error of the mean. Solid lines show best fit to changes with eccentricity, dashed lines are bootstrap 95% confidence intervals determined by bootstrapping. **a** Model fit variance explained as a function of preferred visual field position eccentricity for visual field maps in early visual cortex (V1-V3) and lateral-occipital (LO1-LO2) visual field maps for participants P1-P5 with Fourier power model data. **b** Visual field maps in temporal-occipital (TO1-TO2), parietal association (V3AB, IPS0-IPS5) and frontal (sPCS1-sPCS2, iPCS) areas for all participants. **c** Progression of monotonically increasing Fourier power and monotonically increasing numerosity response model fits with eccentricity was best captured by a cumulative Gaussian sigmoid function in V1–V3 and LO1 visual field maps. For all other maps and models, the eccentricity progression was better fit by a quadratic function. Solid and dashed lines show the goodness of fit of the cumulative Gaussian and quadratic functions to the eccentricity progression, respectively.

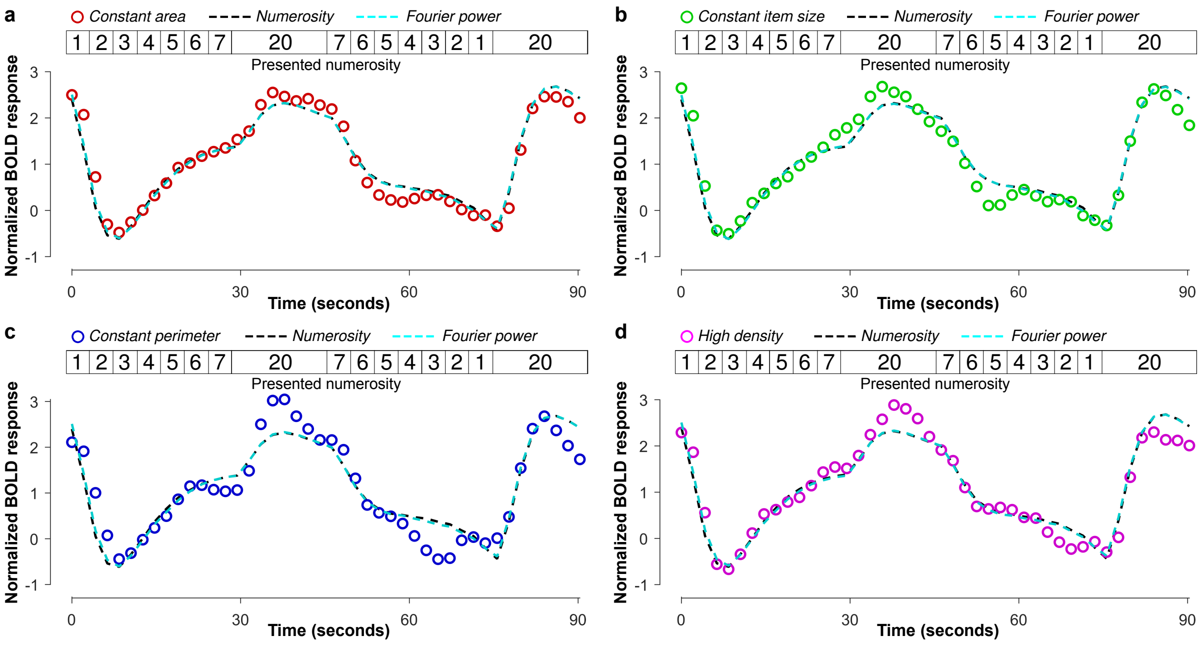


**Supplementary Fig. 6:** Monotonic brain responses across the seven numerosities (and baseline numerosity 20) for each configuration together with numerosity and aggregate Fourier power model predictions for the same conditions. **a** Constant area. **b** Constant item size. **c** Constant perimeter. **d** High density.

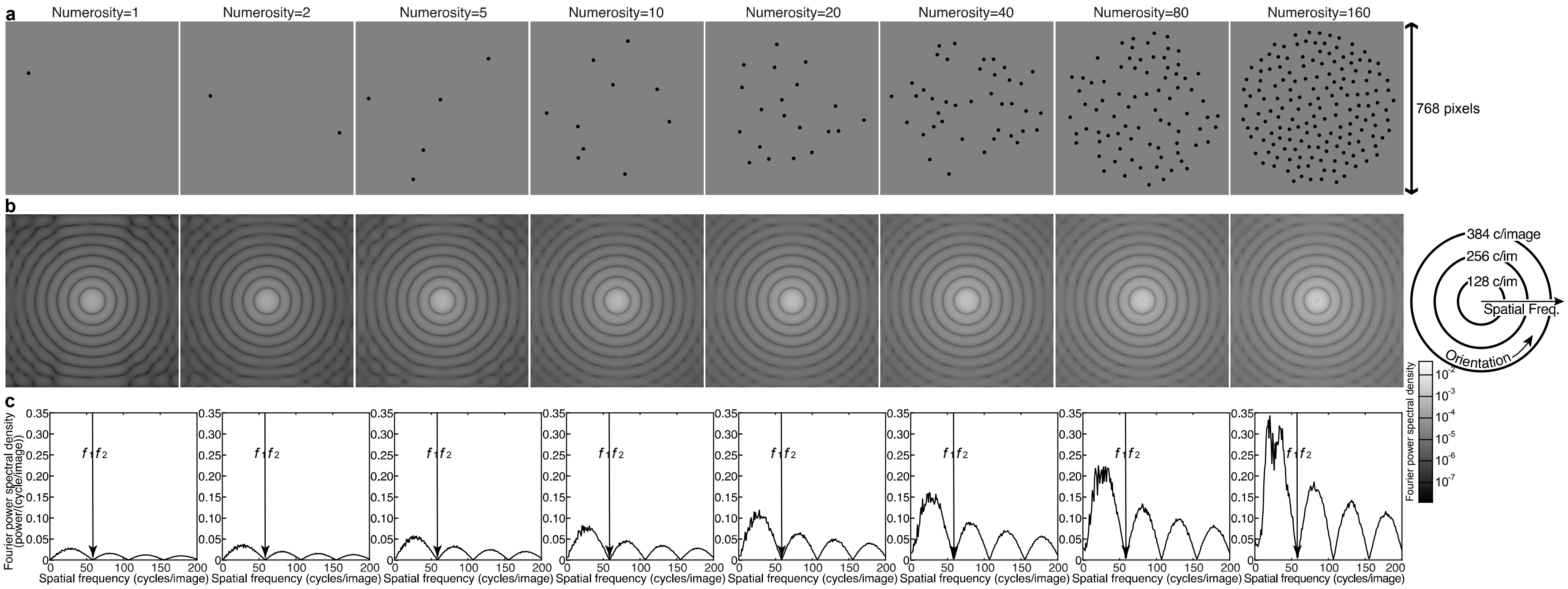


**Supplementary Fig. 7: Fourier power spectra and display numerosity.** **a** Randomly generated images with a parametric increase in numerosity from 1 to 160 items, using the same iterative procedure used to draw the stimulus configurations shown in our experiment (constant item size, non-overlapping, sampled from a central circular area). **b** Fourier transforms of each corresponding stimulus image shown in panel **a**. **c** Fourier power spectral density each spatial frequency, collapsed over orientation, of the corresponding transformed images shown in **b**. The limit of the first harmonic ($f_1$) is used to determine the aggregate Fourier power of each image.

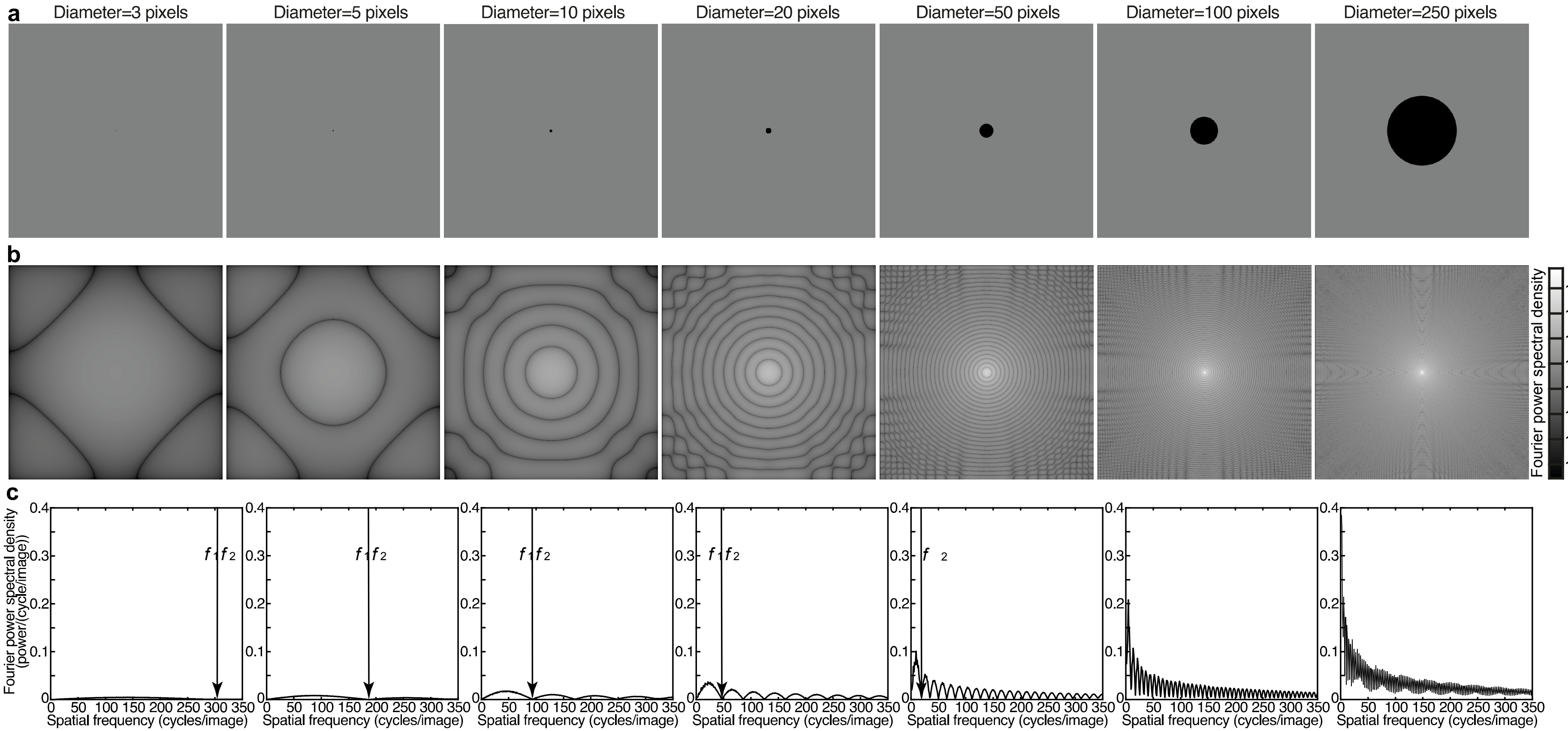


**Supplementary Fig. 8: Fourier power spectra and item size. a** Illustrative images of a parametric increase in item diameter for a single circle placed at the image center ranging from 3 to 250 pixels. **b** Fourier transforms of each corresponding stimulus image shown in panel **a**. **c** Fourier power spectral density each spatial frequency, collapsed over orientation, of the corresponding transformed images shown in **b**. The limit of the first harmonic ($f_1$) is used to determine the aggregate Fourier power of each image.

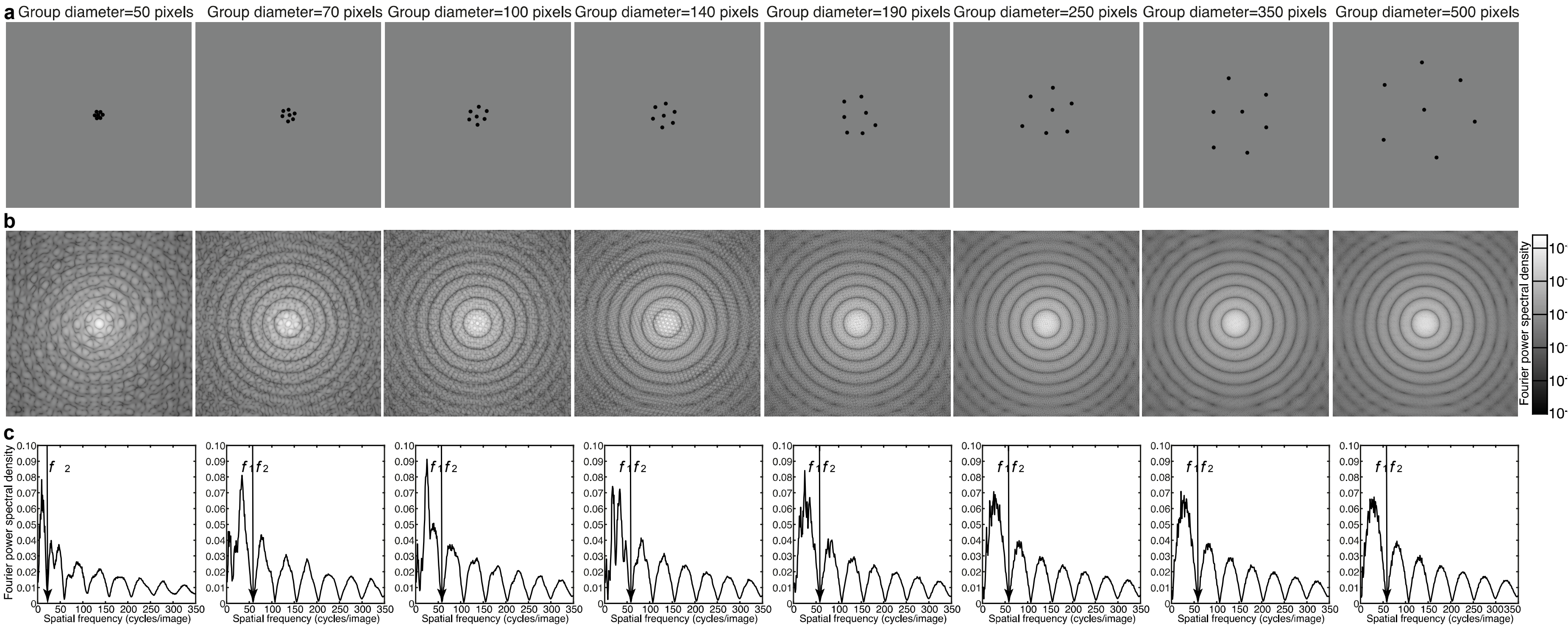


**Supplementary Fig. 9: Fourier power spectra and item spacing. a** Illustrative images of a parametric increase in item spacing for 7 circles, each of 16 pixels diameter, placed at the image center ranging from with changes in the diameter of the stimulus area from 50 to 500 pixels. **b** Fourier transforms of each corresponding stimulus image shown in panel **a**. **c** Fourier power spectral density each spatial frequency, collapsed over orientation, of the corresponding transformed images shown in **b**. The limit of the first harmonic ($f_1$) is used to determine the aggregate Fourier power of each image.

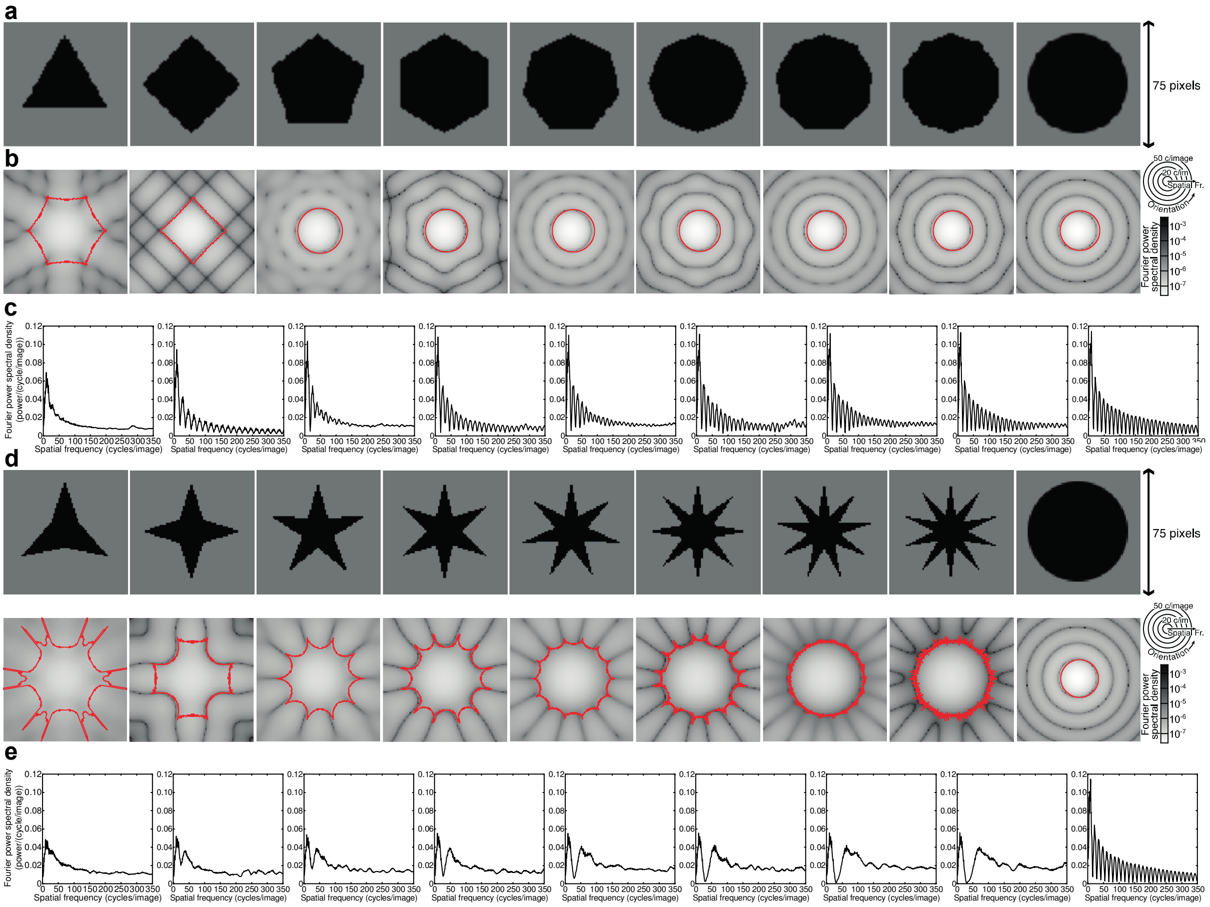


**Supplementary Fig. 10: Fourier power spectra and item shape. a** Regular polygons with a parametric increase in the number of corners from 3 to 10, as well as a circle (infinite corners). **b** Fourier transforms of each corresponding stimulus image shown in panel **a**, above. The red line shows the limit of the first harmonic at each orientation, and we determined the aggregate Fourier power within that limit. **c** Fourier power spectral density each spatial frequency, collapsed over orientation, of the corresponding transformed images shown in **b**. The limit of the first harmonic differs by orientation, so is not at a single spatial frequency. **d** Stars with a parametric increase in the number of points from 3 to 10, as well as a circle (infinite points). **e** Fourier transforms of each corresponding stimulus image shown in panel **c**, above. The red line shows the limit of the first harmonic at each orientation, and we determined the aggregate Fourier power within that limit. **f** Fourier power spectral density each spatial frequency, collapsed over orientation, of the corresponding transformed images shown in **e**. The limit of the first harmonic differs by orientation, so is not at a single spatial frequency.

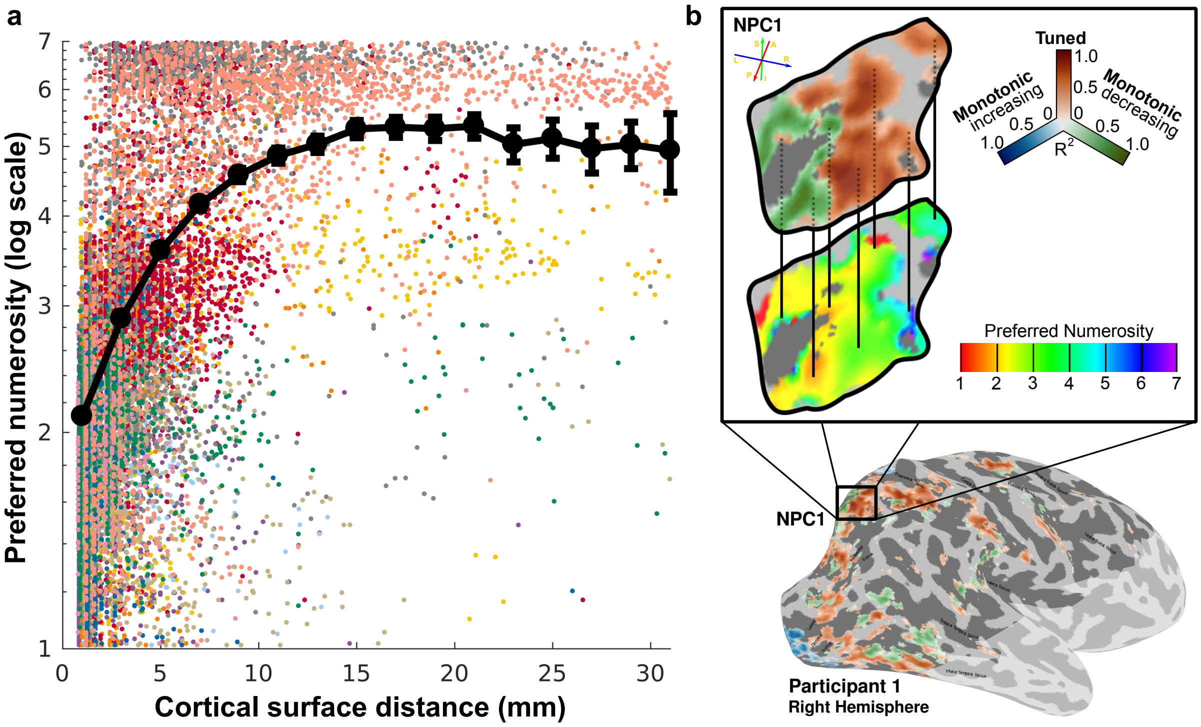


**Supplementary Fig. 11: Monotonically decreasing responses outside early visual cortex.** **a** Preferred numerosity of the nearest recording site to a monotonically decreasing response as a function of distance along the cortical surface (recording sites are organized into 2mm bins; black circles show group mean ± 2 × standard error). Individual points represent recording sites color-coded by participant. **b** Illustrative example of relationship between tuned numerosity preferences and monotonically decreasing responses across cortical distance. Selected region-of-interest (inset) is one of the previously identified topographic maps of tuned numerosity preferences in the right hemisphere of Participant 1; NPC1 (1). Vertical black lines highlight the correspondence between lower tuned numerosity preferences occurring closer to recording sites identified as better fit as monotonically decreasing, and vice versa.

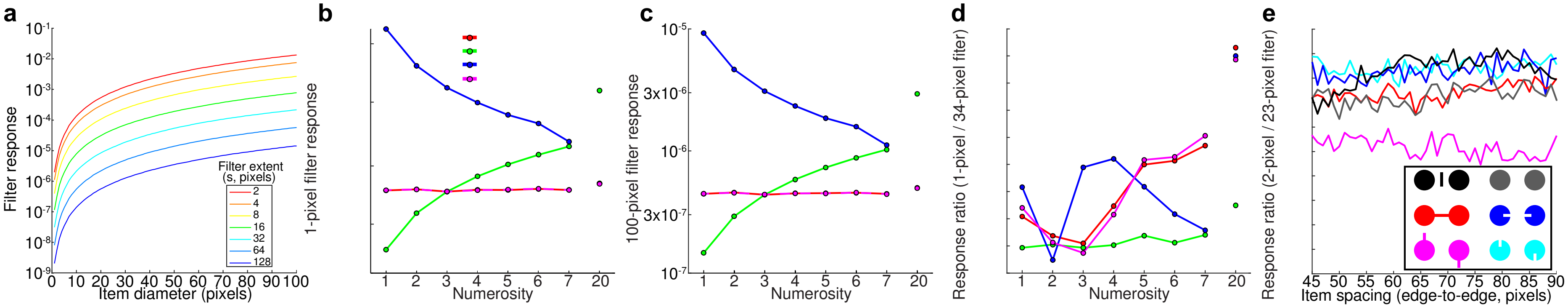


**Supplementary Fig. 12: a** Responses of Dakin and colleagues' high spatial frequency filter response metric to a single circle of different sizes, for a large range of filter extents. Regardless of filter extent, the filters responses increases monotonically with circle size, so no filter of this type can give a response proportional to numerosity if item size changes with numerosity. **b, c** Responses of Dakin and colleagues' high spatial frequency filter response metric to different numerosities in each of our stimulus configurations. This is proportional to numerosity when item size is fixed for all numerosities (as Dakin and colleagues show), but this relationship does not generalize to other stimulus configurations. This is because this metric does not follow numerosity regardless of item size (**a**). The same result is found whether the filter extent is 1 pixel, 100 pixels, or any value in between. Note the red and magenta points and lines lie on top of each other. **d** Responses of Dakin and colleagues' response ratio metric, which they propose subjects actually perceive, for the pair of filter extents giving the highest correlation to numerosity. This metric does not closely follow numerosity, although subjects accurately and spontaneously perceive numerosity in these stimuli with minimal effects of object size and spacing.
